# Novel, visual-attracting traps, accompanying technology, and related behavior of overwintering brown marmorated stink bugs (*Halyomorpha halys* Stal)

**DOI:** 10.64898/2026.06.19.733468

**Authors:** Russell F. Mizell

## Abstract

Objectives identified as critical for effective management of brown marmorated stink bugs, *Halyomorpha halys* Stal (BMSB), were addressed in the overwintering life stage. Research was conducted on a 675-acre farm in western Maryland, U.S.A. Developing and testing were done during September – October of each of the years 2013 – 2020. Traps and related materials were developed. Four invented traps plus the development and use of “weed beater” polyethylene plastic in combination with the other traps were used as attractions to BMSB during the overwintering testing periods. Overwintering BMSB showed very little interest other than to find a place to hide that was dry, dark, easy to get into and safe over the winter months (Nielson et al. 2016). Placement of traps show effect of cardinal directions due to heat and light were north = east > west = south of numbers of BMSB. These data were affected by the vegetation, type of buildings such as silos, cabins, houses, etc., around BMSB prior to the en masse flight. Silos were used as attractants from long range during the en masse flight. Five trap types were invented and tested for use against overwintering BMSB found various places such as buildings of any type, pathways in the woods and other habitats present. Trap #1 was made of plastic out of special plant pots, turned upside down with a plastic lid and entry holes top and bottom that allowed the BMSB to reach the hexcel inside and stay there. Trap #2 was made of white, rectangular coroplast and pressed into 0.5 × 0.5 × 7.32 m (5” × 5” × 24”) box also holding hexcel. Traps #1 and #2 are useful for attachment to most anything with BMSB such as the eaves in houses, other buildings, vegetation, crevasses, etc. They can be placed as horizontal or vertical with other ways that enhance efficacy. Both #1 and #2 are cheap and easy to make and use. Trap #3 is made of a cardboard tube, as a fake or “faux” tree (dead and/or dying) in vegetation 20 cm (6-8”) diameter and 1.22 m (3 or 4’) long. The tubes are placed on tomato stakes, have hexcel and holes made as do the others and are placed vertically in the ground. The tubes are covered with small widths of white, black and or burlap around them. Lids of plastic are on top and the trap appears as a dead and/or dying tree. The fourth trap is a “mechanism” or resembling some kind of “Haven” if you will, made of “weed beater” plastic materials 1.98 m (6’6”) wide in brown or black. The materials are placed on trees by wrapping them around 3-4 trees near 4.6 m (10-15’ long). Traps 1 and 2 can then be placed on the outside of the plastic walls and the #3 fake trees can be added around the outside areas. Around 60,000 BMSB were collected and counted in these experiments by the author, and several different factors were raised and addressed to better understand the BMSB behavior during the overwintering stage. The results provide several new tools as well as understanding of approaches to address BMSB management better.

## Introduction

The brown marmorated stink bug (BMSB) (*Halyomorpha halys* Stal, Hemiptera: Pentatomidae), was first found in the U.S. in the late 1990’s in Pennsylvania and spread early into and around New Jersey. Since then, the insects have spread over a wide host range of agriculture and crops, including tree fruits, small fruits, vegetables and row crops. BMSB’s life history has been well described (Nielsen and Hamilton, 2009). An effort was targeted to better understand the overwintering behavior by developing traps to provide and use for IPM efforts. Watanabe et al. (1994) first studied the overwintering behavior and built “slit” traps developed in Japan to enable reduction of BMSB populations.

Given the early importance of the BMSB as an agricultural pest, a great deal of interest and funded research was conducted to improve management. As with any such situation, starting questions begin and new ones come along based on the results. Because of the need for certain questions to be answered first, as usual, there is time and subject matter picked in that way. Overwintering behavior in general is not that commonly important, so understanding the factors is required to be done in the field over the times they occur. In this case one of the areas that require specific and important information about what happens is learning about the other factors during the year before fall and winter. Developing and testing certain traps effectively are scaled largely into 2 spaces of the year: those for when the insect is active and enabling entry into how they behave over the early portion of the year and following what changes when days get shorter and temperatures colder. Populations are highest late in the season coinciding with the ripening and harvest of many crops (Nielsen and Hamilton, 2009). Venugopal at al. (2015 a, b) provided a well-studied overview of the summer behavior and importance of understanding how BMSB operate in seasonal spatiotemporal soybean fields, edge effects in plant nurseries and in tree borders along with patterns of abundance in such fields and habitat types. Gathering field results from these efforts provided information on field edges of 5 m and 15-20 m away from edges and indicated that BMSB nymphs and adults were more abundant on edges. Soybean fields and nurseries in general were used in high numbers and the samplings indicated that specific areas related to the field edges might best be the more specific places to use for management needs. In another evaluation of the BMSB along tree borders and other host trees, for example, the “tree of heaven” (*Ailanthus altissima* Mill.) along with soybeans, it was found (Aigner et al. 2017) that BMSD built up the numbers on the *A. altissima* trees on the soybean borders in July then moved toward the soybean edges later. Two field edges containing soybeans were evaluated at 15 and 30 m with most staying within the 15 m range. These results indicate that monitoring and management efforts should be most important in areas closer to the edges and trees. The behavior of BMSB in the nymph and adult stage changes as the daylength changes due to overwintering (Leskey and Neilson 2018). Unfortunately, the summer behaviors of BMSB found in fields continues along with the change to overwintering behavior.

Understanding behavior, time and movement of the first summer sections require different traps and lures and will not be discussed here (Bergh et al. 2017, Rice et al. 2017, Possebom and Reisig 2016). Bergh et al. (2017) worked on spring emergence, wooden shelters and pyramid traps and found that emergence was early in mid-April then in larger numbers by mid-May and June. There results also indicated the likely occurrence of a dispersal flight before the response to pheromonal-baited traps. Developing and destroying crops by BMSB are the first very important areas and overwintering information follows later. Of course, entomologists have many standard ways to ask and answer the required questions to move forward. Leskey and Nielson (2018) discussed and summed up what was known up to year 2018. Related overwintering factors will be adding the required and available information that were gathered and used effectively to address how the BMSB overwinters and how questions could be set up and used as an example testing of newly made traps.

BMSB is unique among stink bug species in that, adults commonly overwinter in human-made structures (including homes, sheds, silos, barns, etc.) as well as in sheltered sites in wooded areas or cliff outcroppings, etc. Watanabe et al. (1994). The nuisance factor associated with overwintering populations garnered significant public attention before it was recognized as an agricultural pest. In fact, Inkley (2012) reported on the number of BMSB that were found over 181 collection days as 26,205 in one home. BMSB are so well known by now that there are a number of people who built and now provide various methods to remove the BMSB from inside households provided on YouTude™. Adult BMSB from the first generation can live until midsummer and feed and reproduce throughout that period. As a result, the overwintering (P1 generation) is capable of producing 2nd and some 3^rd^ generations per year depending on locations. In early autumn, adults begin to disperse to overwintering sites. We will identify characteristics of human-made overwintering sites that will complement identifying BMSB overwintering site selection within natural landscapes. Information generated regarding mobility and timing is supported by Nielsen et al. (2016, 2017). Whole-farm BMSB management of BMSB life stages and movement patterns at farmscapes level will complement landscape-oriented objectives.

Adult BMSB overwinter in human-made and natural structures, with peak dispersal to overwintering sites occurring during fall days such as in September (e.g., Maryland) depending at what north or south areas they live and emerging from these sites in Spring of the following year. To determine an optimal overwintering trapping ability, combined data were collected and tested from natural overwintering sites and from year 1 of identifying common characteristics - color, shape, size, temperature, relative humidity, etc., of preferred overwintering sites. Trap designs were deployed and trap designs will incorporate overwintering site characteristics. Change this to MD and it changes from each year. This predictable movement provides the opportunity to develop interception traps that serve to mimic preferred overwintering sites and at some point, be used to trap out immigrating and overwintering BMSB fall populations.

Precision agriculture is a relatively new and different type approach to use for IPM on BMSB. There are two European countries, Greece and Italy, that have worked on finding the active seasonal populations using the Normalized Difference Vegetation Index (NDVI) and NDWI (Normalized Difference Index) (Liakos et al. 2023) and the latest satellite information equipment (Giannetti et al. 2024). The data from kiwi orchards in Greece showed that BMSB caught using traps and lures were found where the indexes were highest: more water and at higher altitudes. In Italy, artificial intelligence and drones were used to find high resolution images of both adults and intermediate images. AI models were able to detect with an accuracy of 97%. Description here of the invented BMSB overwintering traps will follow up on this latest technology with respect to use and efficacy. With the newest way to find the BMSB, we may now have the best way to treat them during the overwintering times. Removal of BMSB from overwintering sites could reduce the amount of pest pressure the following spring. Optimal trap design, background information and orientation will be identified.

Objectives are: 1. to develop an in-depth understanding of the behavior and its operational parameters (stimuli-cues, responses) to characterize the behavior into a working descriptive model(s) for testing, 2. to develop and optimize trap(s) capable of reliably quantifying the populations involved, and 3. to test and apply the traps under different scenarios or to answer specific questions as indicated by the model for use in further research and population assessments.

## Methods and materials

### First Year

Research was experimentally focused on the overwintering behavior of BMSB adults and was conducted in western Maryland ∼ 20 miles west of Hagerstown (N 39^0^ 41.160’, W 77^0^ 59.614’, altitude of 192 m) near the town of Big Pool. The experiments were conducted during September – October for each of the years on a 675-acre farm. Early trapping experiments were conducted to determine the behavior surrounding BMSB adult overwintering flights and post flight behaviors. Various materials were used toward the idea of finding the best “trap” like materials. The first materials used were paper, cloth, cardboard pieces and the like. At that time “hexcel” materials were found to work best and applied as well. One trap was placed in each cardinal direction on the building walls ∼at 1.5 m above ground and ∼0.5 m below the building roof edges. Buildings one and two were utility sheds each 3.1 × 3.7 m in width and length, respectively. Building there was a log cabin ∼3.7 × 4.9 m. Two interior traps were placed in building one on the north and west walls. Numbers of BMSB were much lower in 2014 than in 2013 at this location. All of this has the ultimate goals of development and implementation of new suppression strategies and tactics.

#### Methods

A working descriptive model of BMSB overwintering behavior was deduced from previous research (PI and others) and tested as appropriate from silos, barns, trees, houses and cabin buildings. Several trap parameters including size, color, configuration, orientation, context and placement relative to buildings and preliminarily in the landscape relative to structures – corridors, barriers and matrix, etc., were tested (Tewksbury et al. 2003). Note that these factors will be discussed later. This was executed by placing different types of traps or sets of traps in comparative side by side or replicated treatments on various types of buildings or other structures: houses, adjacent sheds, barns, silos and adjacent feeding structures – out buildings, as well as in the surrounding farm and forest landscapes. Pictures of these areas and how they were used are provided in the figure sections. Silo alone without any type of materials that would be used by BMSB are only of use driving the yearly en masse flights. The overwintering BMSB then moves from the large silos to other materials that are of BMSB use unless there are materials attached to the silos.

#### Results

Less than 3% of the ∼ 225+ original traps in the first year used did not capture at least one stink bug while trap counts were as high as 500+ in others. BMSB populations varied greatly among the buildings used and even more so in the landscapes sampled. However, ∼9,000 BMSB adults were counted during this study (year 1) by the author. The behavior of BMSB in response to structures and traps around or on them is highly influenced and confounded by their macro-and micro-habitat context. Moreover, the behavior in response to cues during arrival flight is not always the same (or at least indicative of) the local behaviors on structures after arrival in response to traps. See the personal drawing of BMSB’s potential biology and ecology behavior (Fig. 1) that the author developed along with other literature. Thus, the individual trap captures are the end of a long behavioral sequence and caution is warranted in interpretation of results. Once as an example early in this work, a clear plastic sheet 9 m × 6.6 m (30’ × 6’) long and wide with lots of different black painted forms was covered with Tangle Foot™ (sticky material) and placed up on the southwest side of a silo and overwintering BMSB completely covered the sheet by the 1,000’s.

**Fig 1:**
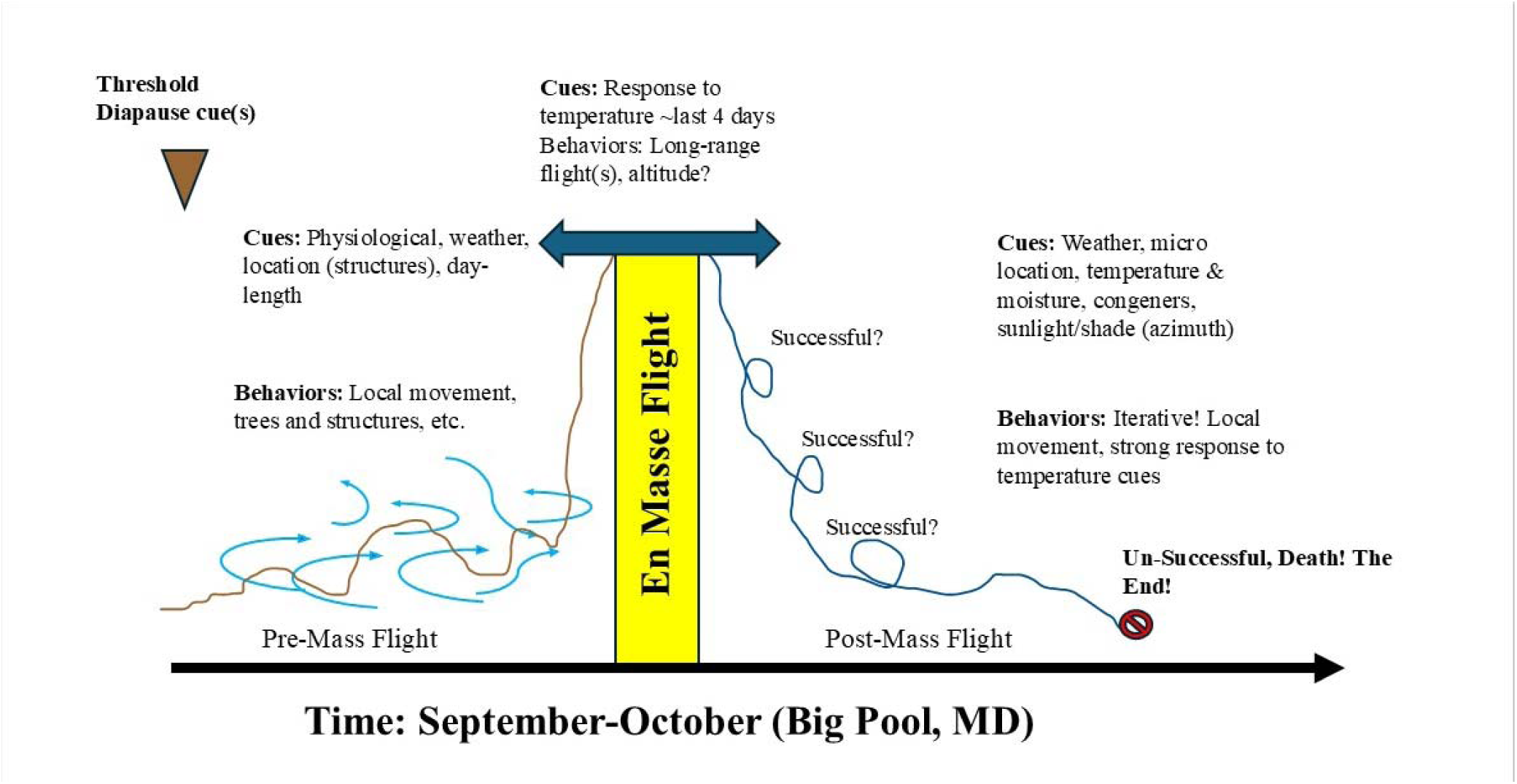
En masse flight pattern made by the author: Brown marmorated stink bugs (BMSB) are unusual doing the overwintering diapause period. In the fall they look for places to hide over the winter. They hide in dead and dying trees and all kinds of other places (see within), but if they cannot find what they need, they respond to the weather and search using an en masse flight together. This figure explains what happens to them during that activity.

Thus, no statistical differences were observed among treatments, but numerically, a number of trends were manifested by the data as follows with all counts equal to mean ± SE number of overwintering BMSB adults. Trap color: white 117 ± 26, cardboard tan 79 ± 37, yellow 78 ± 26 and silver 32 ± 8. Trap size: “standard” plant pot 11.35L (3 gal, 770 cm^2^ surface area) 77 ± 33, plant pot 11.35L (3 gal, 760 cm^2^ surface area, scalloped sided with extra holes) 27 ± 8, plant pot 12.9L (3 gal, 866 cm^2^ surface area with striations) 43 ± 17, plant pot 27L (6 gal, 980 cm^2^ surface area) 46 ± 14, plant pot 34L (9 gal, 1160 cm^2^ surface area) 63 ± 16. Trap height using the “standard” plant pot above was affixed to a silo at different heights. Paired traps were placed but 4 of 16 were blown down due to hurricane winds. Where 2 remained, the counts were averaged (N=4 of 8), otherwise a single trap count is reported. Based on the 4 available trap pairs, there was very little difference between traps at the same height (Fig. 2). This result and similar, prior-year sticky trap collections during the en masse flight, indicate that the BMSB land on the silo after flying from some distance to it at between ground level and heights up to 12.2 m (∼40’), and then took short flights and walks on the silo surface to find the traps. Others responded to structures below and surrounding the silo and its other “feeding” parts and were trapped there (Fig. 3). Note that the traps used were attractive to the insects when they found and entered the traps. Diurnal sequence of BMSB arrival to several buildings (aggregate totals observed on all buildings at each time interval) on 4 warm days prior to the mass flight (Fig. 4). BMSB arrival peaked in the afternoon when warmest temperatures occurred as would be expected. Stickem traps and weird colors show the plastic squares mounted in or on the sides of cabins, homes and barns (Fig. 5) and Fig. 6 is the older and common, plain black pots with the BMSB counts also (Fig 7). The early used cardboard traps of different colors were also tested within the silo augur area. They were also used and tested as simple as possible as re-managed plant pots of different sizes and placed on sheds to determine if BMSB would enter and stay with them. Different types of materials (before using hexcel) were placed in the old traps and evaluated. Things like chopped-up paper, pieces of cotton and other small pieces of materials were used. They did work well enough to consider the traps for more changes. Another test at the bottom of the silo using a silage augur was used and also had a roof and lots of wood braces used to mount cardboard traps which were of different colors – white, yellow and tan cardboard (Fig. 8). The BMSB responded to the areas and were found under and in the traps in low numbers.

**Fig. 2:**
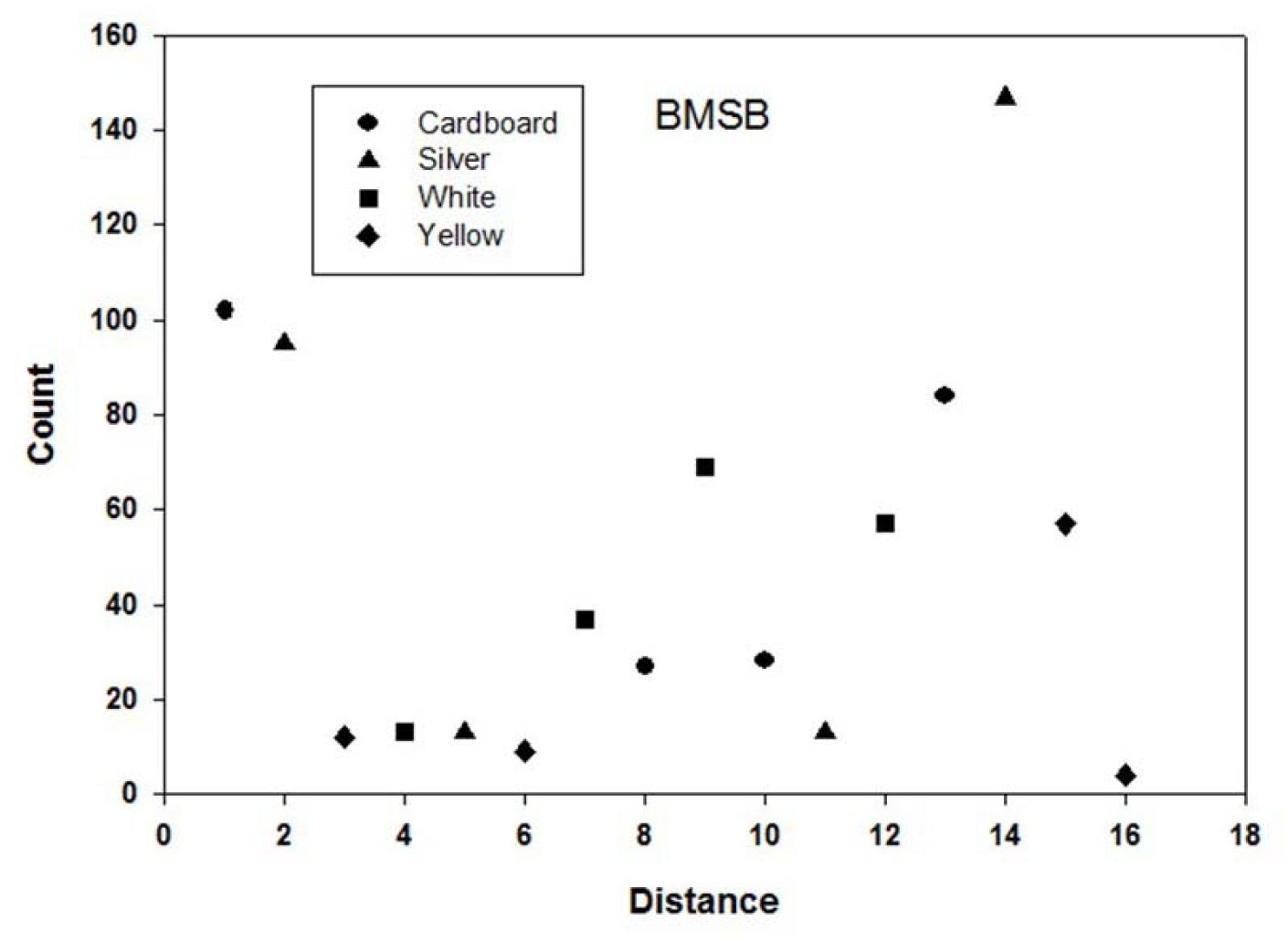
This is the first evaluation of the BMSB overwintering behavior observed to respond to a silo augur holding traps of different colors spread over the distance from the silo base. They moved over the areas (see Fig. 9 for the equipment involved) and move away from the silo and into the silo’s augur spaces along with using all the 4 different colors.

**Fig. 3:**
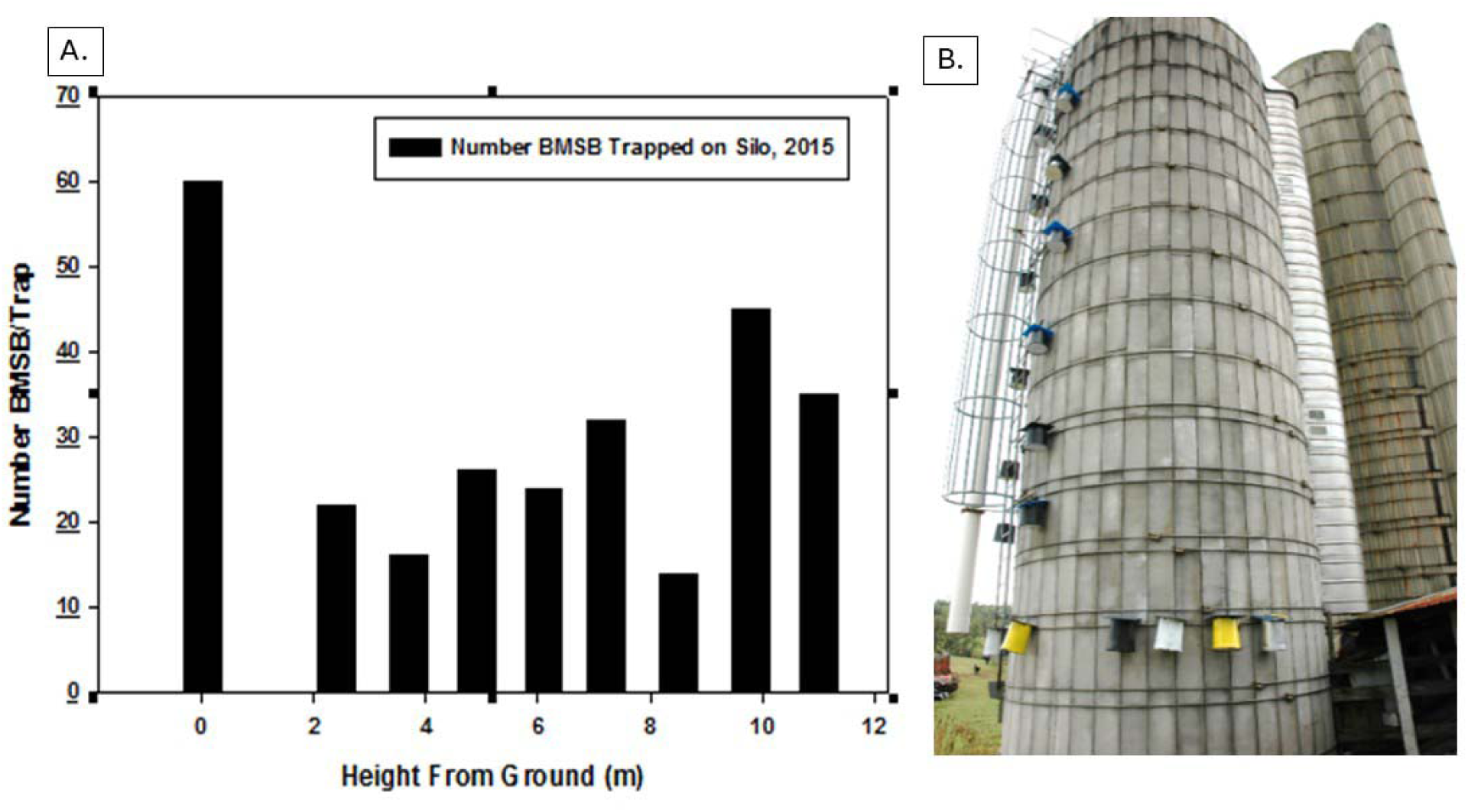
This data was developed by placing traps on the side of a silo ∼40‘ high and bottom area with the 9 traps spaced 8’ apart. There were also colored traps ringing around the silo bottom. The largest number stayed at the bottom area but many chose their traps from all different heights.

**Fig. 4:**
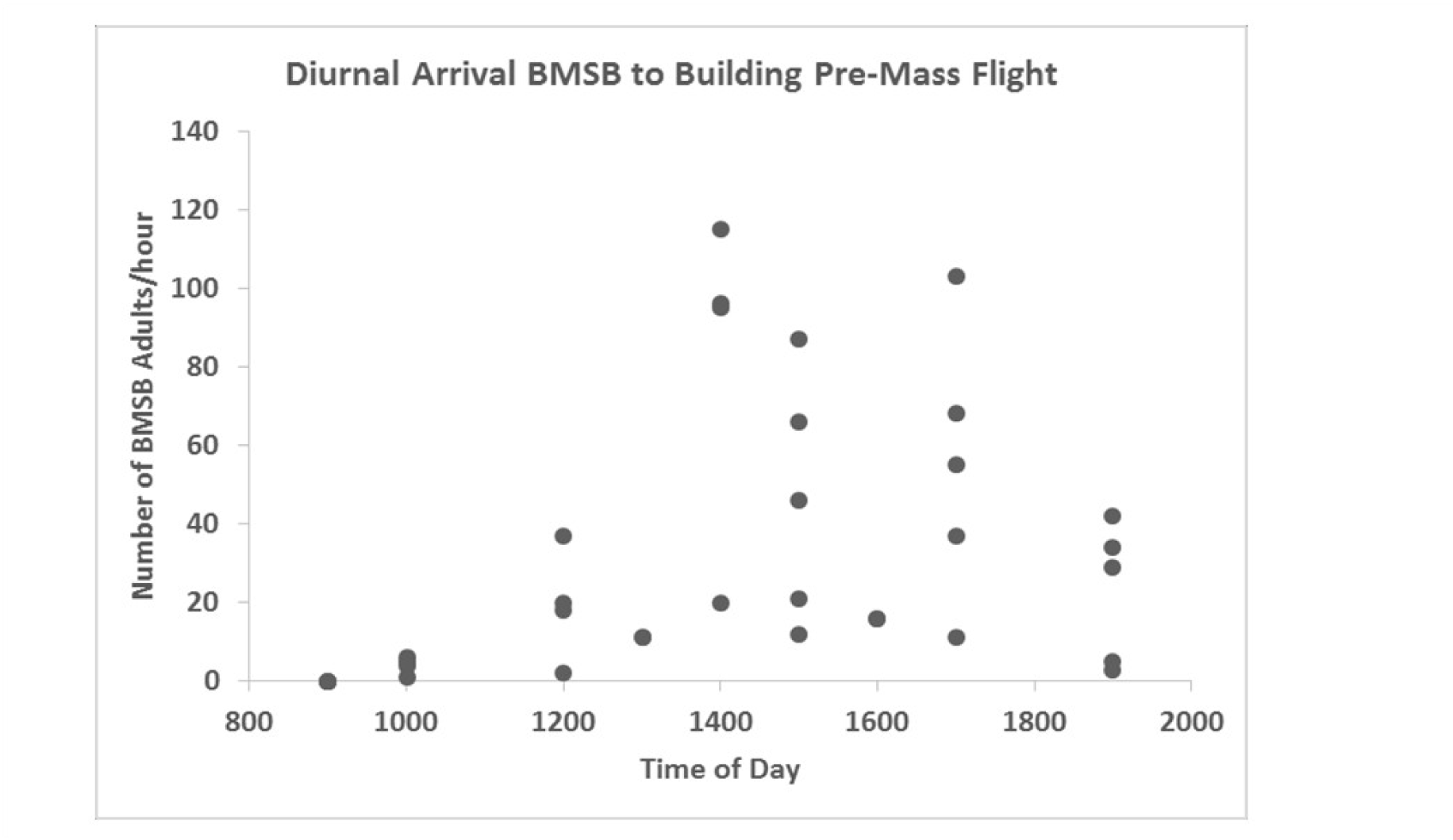
To determine the movement by BMSB through a full day’s time, BMSB on the traps were counted on a two-hour daily pattern for 4 days. The results indicate most of the observed movements by the BMSB occurred in the middle afternoon from 1400-1700. This occurred in early September prior to the en masse flight observed later on.

**Fig. 5:**
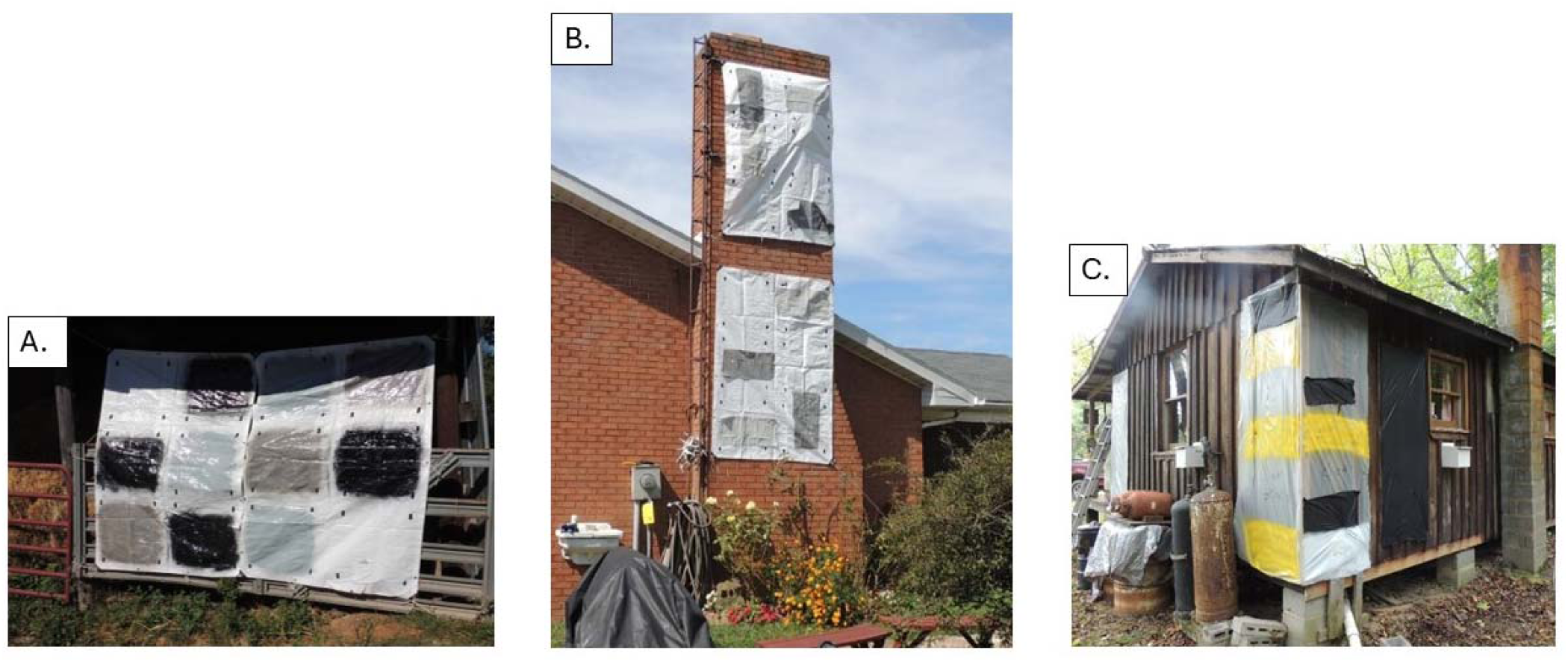
This shows early work with silos and other places on the Mizell farm using TangleFoot™ (sticky material) with various colors and heights. The BMSB did not respond to the traps in any place they were tested.

**Fig. 6:**
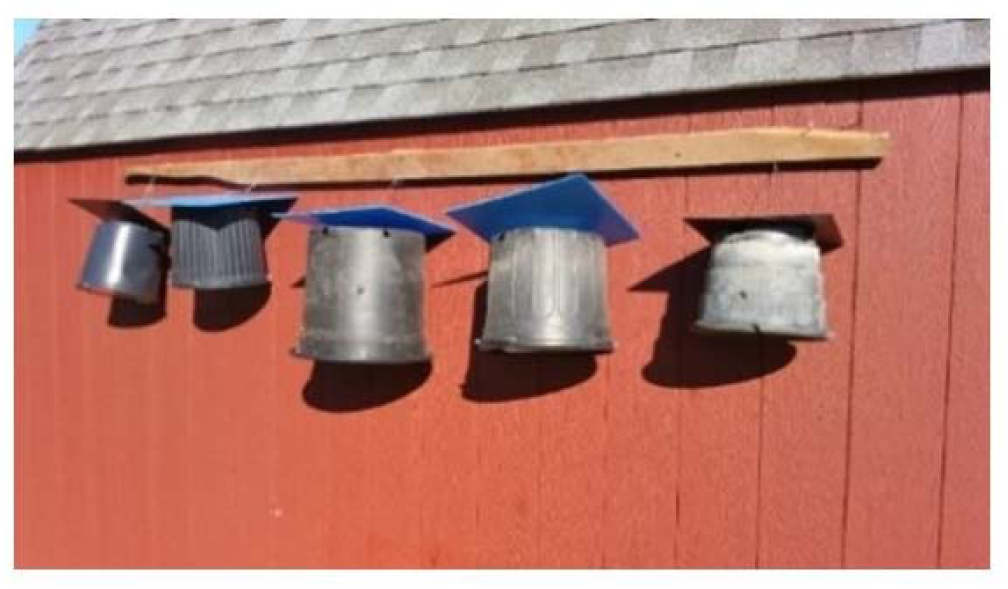
Regular potted plant pots of different sizes were the first tested traps. These traps were turned upside down with an added top, wire handle and bottom placed on a building. The traps also had hexcel inside to capture flying, overwintering BMSB. Numbers are shown on the next figure, Fig. 7.

**Fig. 7:**
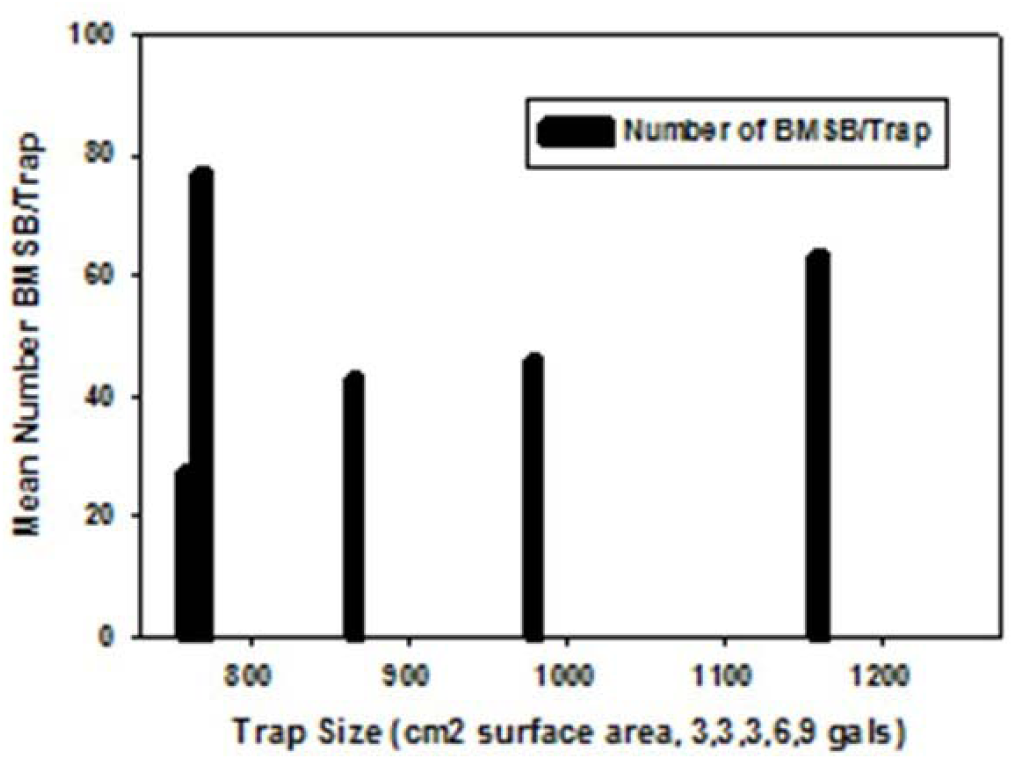
The data from plant pots (Fig. 6) indicate that there were no significant numbers of differences of collected BMSB in the traps of different sizes. The traps were used by the BMSB landing on the building that was behind a house along the woods and further away from the silos.

**Fig. 8:**
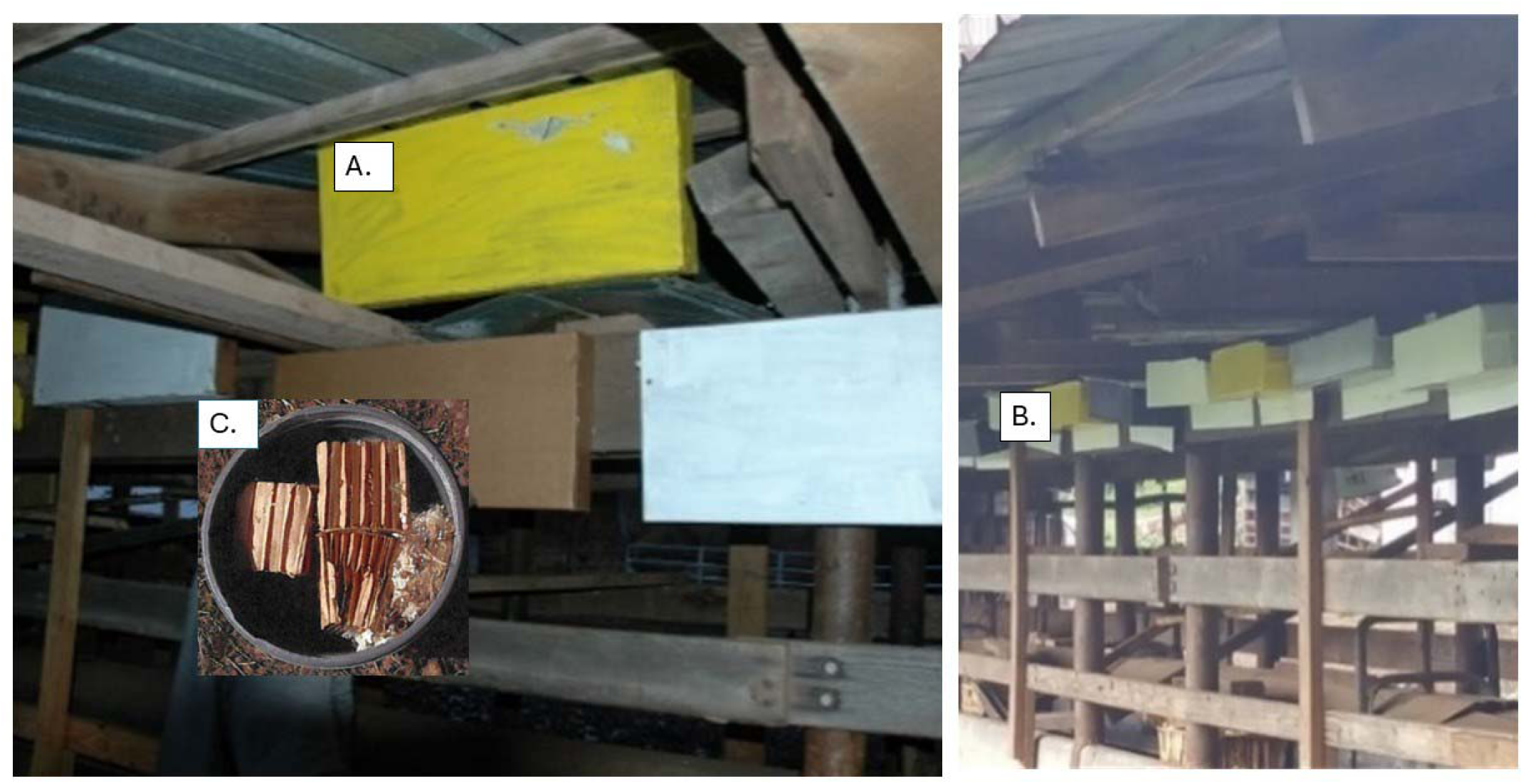
There were plenty of open areas within the silo augur area and medium sized cardboard boxes were provided in white, yellow and tan and tested. Again, there were some BMSB collected but no statistical differences between the numbers of each color. A. one of the early test traps is thin cardboard with cardboard pieces inside it and small holes around the edges. B. shows the coroplast (Trap #2) made trap placed on the wooden tops around the silo augur. The bottom of the augur is shown in the middle and is inside what appears to be a tube. Lots of BMSB were found in this approximately ∼12 m (40 ft) long area. C. shows the 10 by 10 cm (4” × 4”) blocks of hexcel used in the traps.

Silos were available to determine how the BMSB would respond to various heights and several trap positions at different heights. Due to some bad windstorms that year, several of the test traps were lost. The remaining ones showed that more insects responded by staying on the ground with those traps and varied quite a bit in numbers up and down the silo. The silos also were used to determine if the BMSB would respond to a silo from ground to 30 feet high with unusual patterns of black on white with stickem. The BMSB did not respond to any patterns or shapes of black on a white silo (Fig 9) because the number of BMSB was high numbers (thousands) enough to cover all the stickem. Only a few BMSB were found outdoors on these stickem covered traps. With the first year somewhat successful, a larger effort was made to develop, test and use better built and effective traps.

**Fig. 9:**
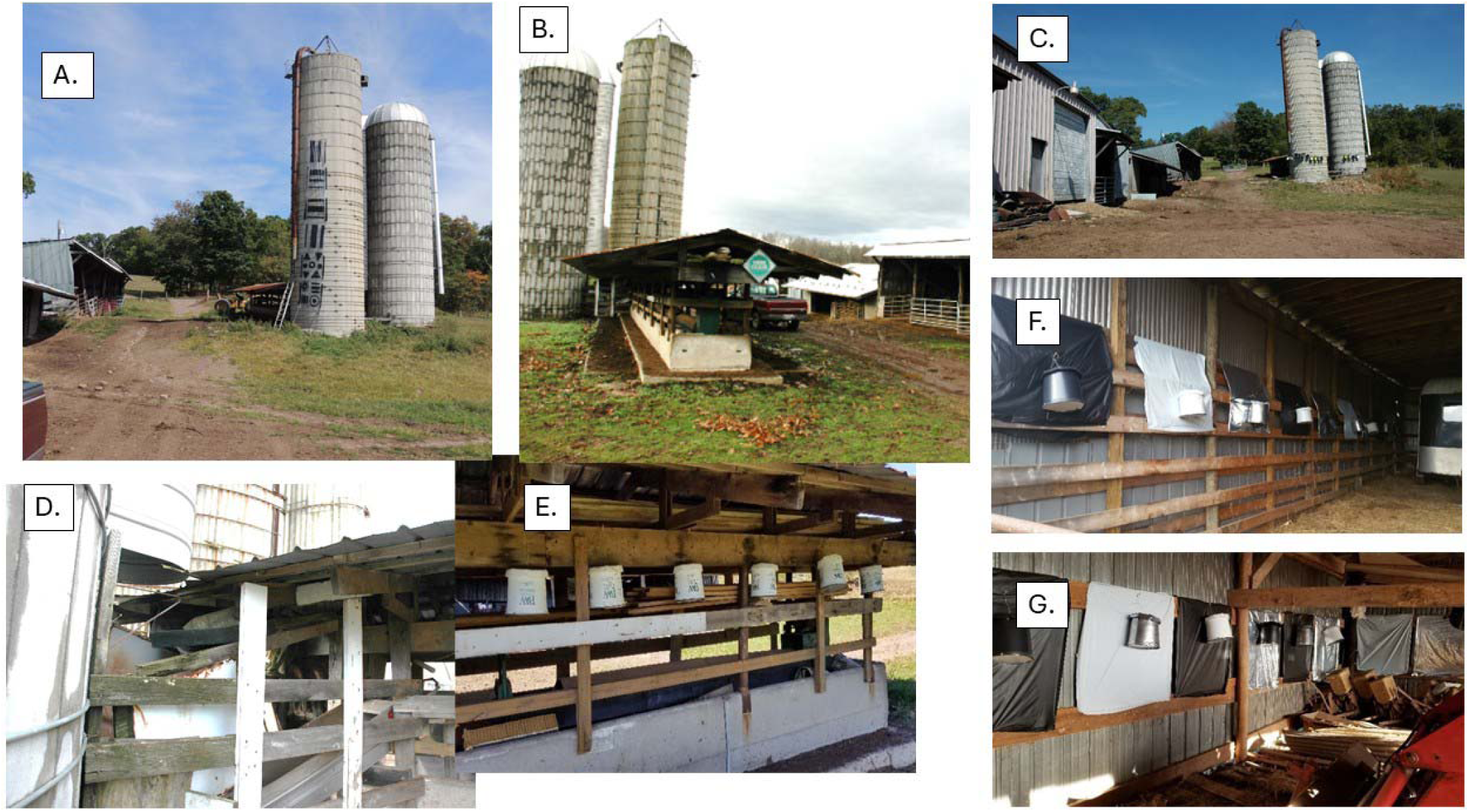
This shows the natural pieces found with a silo of different sizes and covered with parts used on it. In this silo testing of an en masse flight, a large piece of ∼30’ clear plastic. A. was covered using black paint with different patterns and sizes and hung up on the side of the silo. The BMSB did not respond to any patterns or shapes of black on a white silo because the number of BMSB were high numbers (thousands) and enough to cover all the other patterns. B. is the silo augur facing away from the silos. C. is on the other side of the silos showing several different barns. D. shows the base of the silos where a lot of BMSB landed and went into hidden materials around it. E. shows the Trap #1 hanging on the end of the silo augur space. F. and G. are the inside of two of the barns seen inside the C. showing where the trap color vs background colors were tested.

### Following years

BMSB overwintering trap details and behaviors discovered and exploited. Note: during trap developments the goal was to use simple, cheap and readily available materials. I recognize that the designs can be manufactured later in different forms to achieve the same results. Please note that the “key” material used in the traps is what is termed ‘hexcel’ and is made by a leading supply company named hexcel in Washington, Georgia, U.S.A. There are other companies around the world that make and sell similar products such as “honicel”. Also, the company Uline, Inc. carries and sells it. The pieces of hexcel used here were made into 122 × 122 × 10 cm (4” × 4” × 4”} and purchased from a local production company in Rome, GA and used in all traps. Other dimensions are below.

### Invention of effective traps

Five traps and related materials with different structures have been developed via selecting out of many tested prototypes with similar functions and equivalent efficiency, e.g., depending on how and where traps are deployed. The first label is **Trap #1** and termed the “Pot trap” which is cheap, easy to make, use and store. It is made from a 11.4 L (3 gal.) specific plastic plant pot used by ‘Proven Winners™ and made by several companies including the #PP1200 by Nursery Supply, Inc. It is either obtained in white or painted white and turned upside down. This plant pot is somewhat unique in that it has 5 beveled 1.25 cm (½”) holes in the bottom (1 in the center and 4 on the edges) which become the top of the trap and allow BMSB entry under the cap, e.g., top lid. The cap can be handmade out of white 2 or 4 mm coroplast sheeting which is seen in the figures or one can use a 20-25 cm (8 −10”) size clear-plastic, plant water dish used under potted plants that is painted white. A piece of 20 mil wire is attached to a piece of coroplast, wood, whatever (bigger than the center hole) at one end while the other end is then pulled through the center hole and the cap and then used to handle and mount the Pot trap. Trap bottom: the bottom, which was the normal pot opening (at the top), is closed using a round 1.25 cm (1/2”) thick piece of insulation material (light weight) that fits into the pot grooves tightly with slit entry openings on 2-3 sides (partly built into pot and large enough for bugs to squeeze through). These holes are similar to the beveled holes at the top. The bottom is attached to the inside from the outside edge of the pot using two special concrete nails (with plastic collars) making them easily removable (Fig. 10). Inside, the trap contains 2-3 pieces of cardboard hexcel (see https://www.hexcel.com/Products/Honeycomb/HexWeb-Honeycomb) 10 × 10 × 20 cm (∼ 4” × 4” × 8”) or less hexcel cells themselves are 1.2 cm (∼½”) in diameter and diamond shaped. The honeycomb = hexcel pieces are placed inside the pot after the wire handle is inserted and before the bottom lid is attached. The hexcel is made by the company to specs 0.64 – 15 cm (1/4” to 6”) honeycomb cells, etc., with or without both sides covered, e.g., normal use. Note: this is the same type of cardboard material used by biological control producers (using smaller size) for raising predatory insects such as green lacewings, etc. For BMSB traps, the cover from one side is removed (peeled off) and the other side supports the hexcel and provides the large number of holes and the interior dark surface areas for the BMSB to enter and fill. The highest number I ever caught was ∼4,500 BMSB adults in a single trap of this type. The configuration of the pot with beveled entry holes under a lid (critical feature to keep hexcel dry), the hexcel material and the cap are critical factors, related to the behavior of the BMSB when searching for an overwintering place, keeping them dry, cool and in the dark. Once the BMSB go into the hexcel, they rarely leave unless: 1. the location is in the direct sun and it gets too hot, or 2. the temperature is un-seasonally high all around and even then, only a few BMSB are moving, and most go back into the same trap after milling around it. This is mostly related to where they are placed relative to direct sunlight. North and east directions on a dwelling are always better than west or south in terms of higher trap catch (Mizell, personal information, see material and methods of the data). Note: this factor was observed by most research people working with overwintering BMSB. Also, this is not much different behavior from that of the ladybug *Harmonia axyridis* (Pallas) which often overwinters in the same locations where this research was developed (Mizell, personal observation). I have tested various trap colors and white is best (see below). This is likely due to less heat when mounted out in the open but in the shade on buildings color is of less importance. Color becomes a secondary issue except on buildings (post insect landing), but as a behavioral cue when they are arriving, it takes on more importance, see below.

**Fig. 10:**
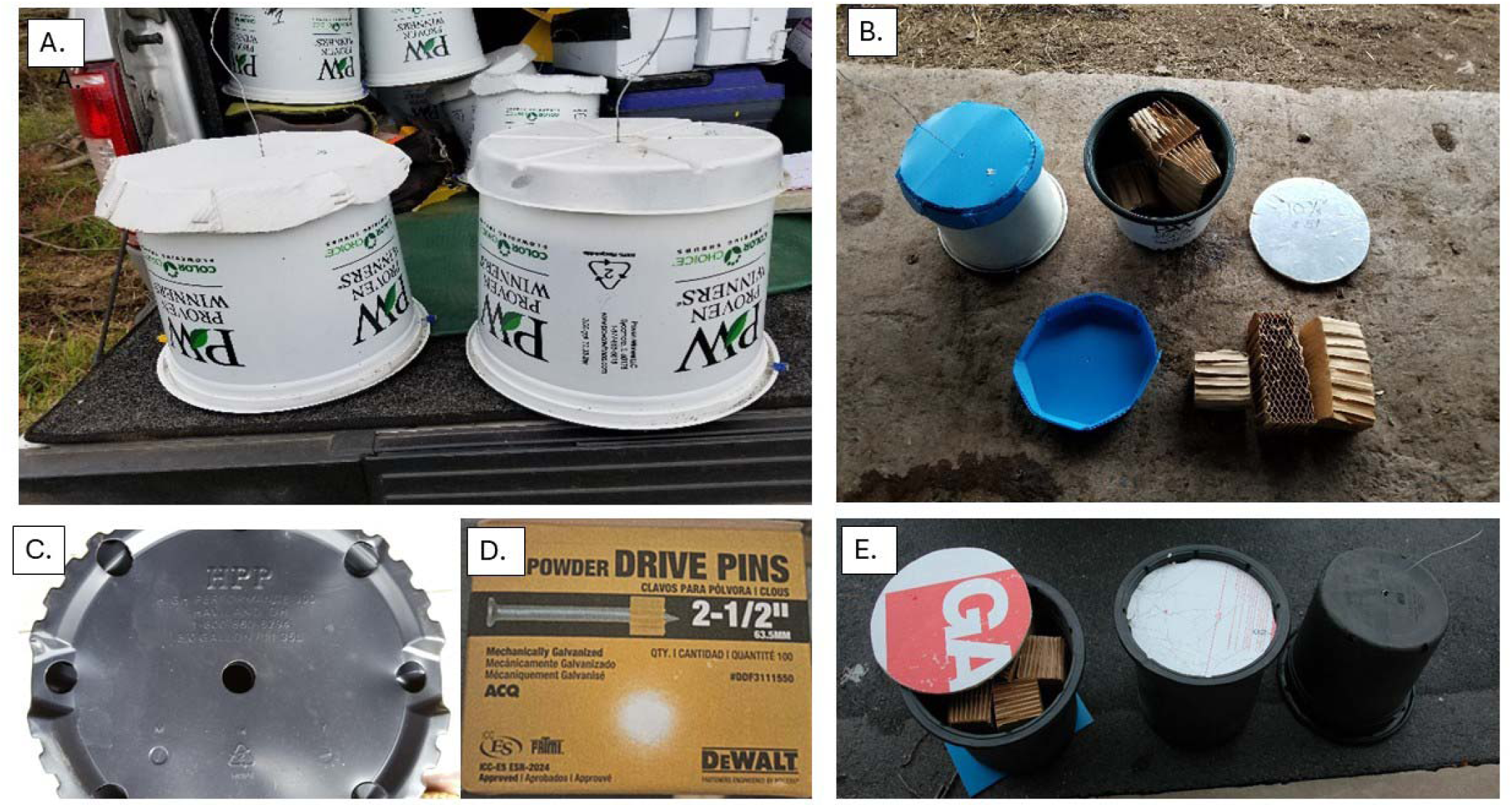
Trap #1 and first invention in this program is presented as the “Pot” trap. It is very similar to common plant pots but is much better built than the regular pots. In this case, the main portion of the trap is the top of the “PW” traps with special slanted holes. The pot is medium size and easy to build or take apart and save for the winter. The trap uses hexcel inside and you can drive pen screws to easily open and close it as is the round cover parts at the bottom. A. shows 2 different tops (one made by hand out of 2 mm coroplast, the other bought) that were used to keep the traps dry outdoors. It also shows a “20 gauge” (20 AWG) wire used in the top of the lids to hang the traps. B. shows all the parts individually. C. shows the special bottom of the PW pots which is turned up where you see the holes with angles that enhance the ability of the BMSb to enter the trap. D. shows the special drive pins that are used (2 each per trap) to hold the round botton part shown in E. next to it.

### Trap 2

The rectangular shaped “coroplast trap” was developed as “easier to use”, with wider uses, and more aesthetically pleasing modification of the Pot trap. The trap is somewhat similar to the pot trap but is larger lengthwise, made from a piece of white 4 mm coroplast material pressed to give the finished dimensions of 12.7 × 12.7 × 61 cm (5 × 5” × 24”) and also contains interiorly the 10 × 10 × 46-61 cm (4” × 4” × 18-24”) hexcel cardboard as does the pot (Fig. 11). Entry by BMSB into the trap is via 1.0-1.3 cm (3/8-1/2”) round holes that are made in the coroplast sides, etc. Entry holes can be customized for how the trap is deployed. The trap can be hung horizontally under eaves and the like or vertically at an apex of the roof and entry holes can be placed strategically to be at the top or sides of the trap that optimizes entry and best keeps rainwater out of the trap, etc. The trap was developed for placement on dwelling walls and works best if placed under the eaves. The trap is light in weight and staples; nails or other means can be used to fasten the trap to building surfaces. The closer the trap is to the top of the dwelling walls (roof eaves) the higher the capture rate and the less exposed it is to the elements. For purposes of using the trap other than on a dwelling, a “lid” of coroplast (as indicated on the Pot trap) can be mounted on the trap out of the same coroplast mentioned with the Pot trap to preclude the entry of rain, etc., if placed on a tree trunk or other surfaces. With this prototype, many holes could be stamped such that the user could later pick which holes are best and customize hole patterns to best deploy the trap for greatest efficiency and rain protection on individual buildings. Again, open holes are not a problem as the BMSB remain (never seen any leaving) in the trap once inside. Both traps can also be used in the landscape (for example, dead-dying trees) and be set up as horizontal or vertical formations.

**Fig. 11:**
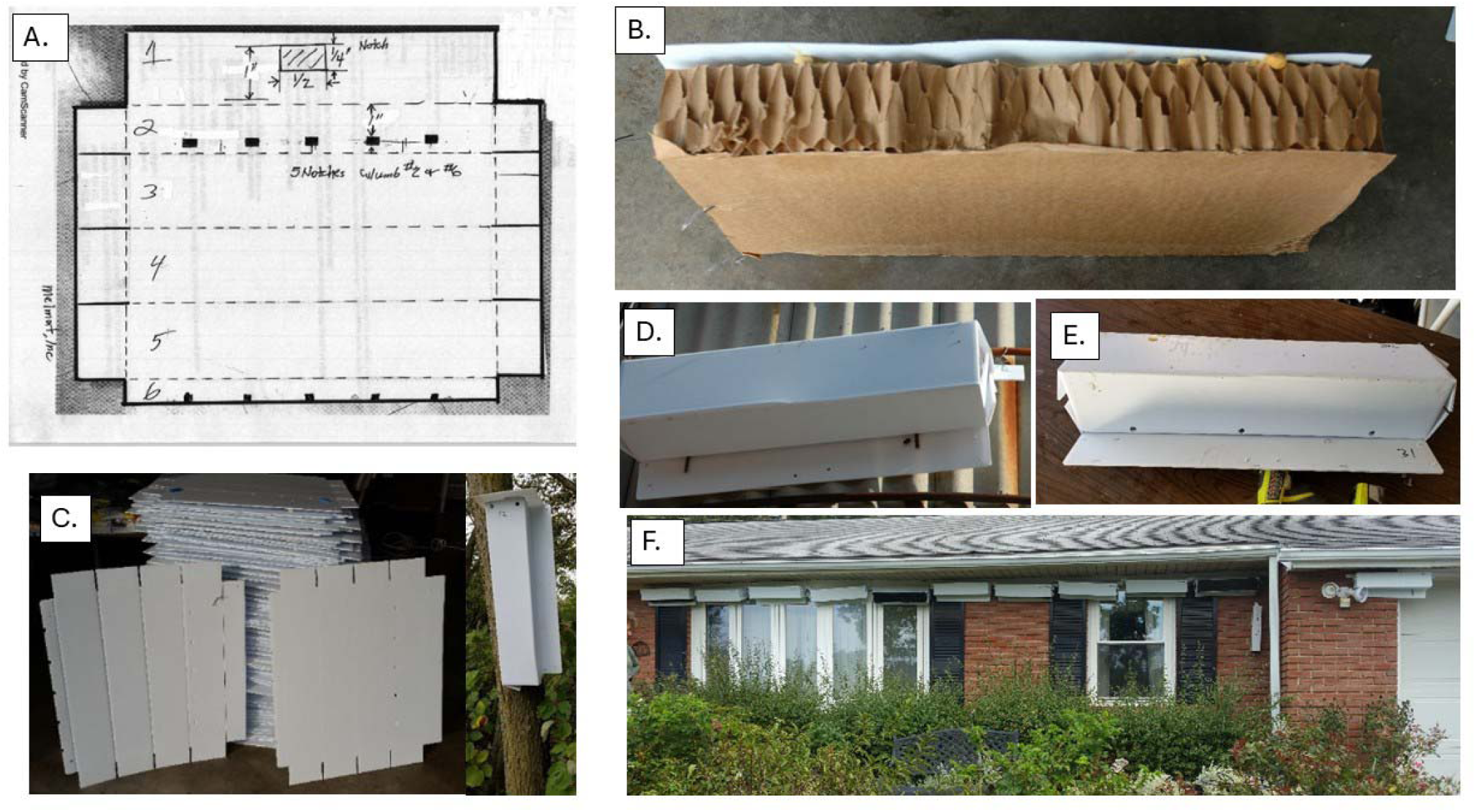
Trap #2 is presented as a rectangular-shaped coroplast trap. The pieces were pressed to make them easy to make and store over the winter. It is easy to put together and has hexcel inside as well as entry holes all around. It can be placed up as a horizontal or vertical shape and lids can be made for each. One of the best uses horizontally is under the eaves of buildings and can be used with the traps at 18-24” apart (different from the picture (F.). A. is the coroplast pressed into shape. B. is a piece of hexcel that can be used as a trap indoors with or without Trap 2 with it. C. shows the pieces used for each trap and the trap hung vertically on a tree. D. and E. show the horizontal traps and F. shows the traps along the eaves of a house before the best size mentioned above was determined.

It was determined experimentally that the traps can be placed >46 cm - 61 cm (18” to ∼24”) (linearly) apart on a wall to maximize sampling space, manipulate BMSB and minimize total trap numbers used on a per linear foot of wall expanse. Example: one 61 cm, (2’) long trap provides 1.8 m (∼6’) of sampled space 61 cm (2’) trap 61 cm (+ 2’) on either side of the trap. Also, the ends are flattened toward the inside of the ends. This enables calculation of the optimum number of traps/wall length. If they are placed within <46 cm – 61 cm (<18-24”) apart, the active sampling space of the traps overlaps and compete, unnecessarily and ineffectively. Again, more BMSBs will be captured on north and east facing walls due to the response of BMSB to lower sunlight and heat (Mizell, personal communication along with many others).

However, the response to building walls is mitigated by the buildings’ surroundings and BMSB do land and search on walls facing other cardinal directions. BMSB mills around buildings and lands a few places depending on the characteristics of the walls; several variables are involved in this behavior. Again, both described prototype traps can be used on dwellings and in the landscapes, as they are or with other configuration materials.

### Trap 3

The third trap is a “Fake” or “Faux” tree mounted on a tomato stake, also placed around buildings, etc., wherever the BMSB inhabit (Fig. 12). Cardboard tubes are used for making concrete building footers and Home Depot and Lowe’s, Inc. among others sell them at 122 cm 20-24 cm (3-4’ long and 8-12”) wide. Boxes provided by Uline, Inc. which are of square shape can also be used with plastic lids, holes in them and hexcel added inside. The use of a tomato stake to mount it is similar to the tubes or Uline boxes 11 m × 2.4 m (36’ × 8’ × 8’), number S-18928, ∼$50.00/15 boxes). Also, coating these types of traps with Flex Seal, Thompson water seal and the like are useful to protect the materials when used outdoors. White materials along with brown colored cardboard tubes and burlap materials affect the attraction of the BMSB during overwintering movements. While the tubes used are larger, they also can be used as standing and appearing as dying or dead trees. Both white plastic materials along with brown or black, - colored cardboard tubes and burlap (painted or cloth materials) affect the attraction of the BMSB during overwintering movements. Small hanging pieces in ¼, 1/3 and ½ size pieces dropping down from top to bottom can help make the fake traps look like dead and or dying trees (Lee and Leskey 2015). I have caught anywhere from 0-100’s to 500’s BMSBs in these individual traps. Burlap fabric is made of brown-tan material readily available in different sizes and costs. Burlap paint can also be used in a smaller part on the tubes with plastic materials such as those used in a tomato field. See Uline, Inc. for an example, S-15521 for 102 cm × 91.4 m (40” × 100 yards) roll. Burlap fabrics and burlap paint were tested to enhance the “faux” trees and this was a success. Using LTM the field results were 15.5, Pr < F = 0.010. Black plus burlap increased results to 0.0087.

**Fig. 12:**
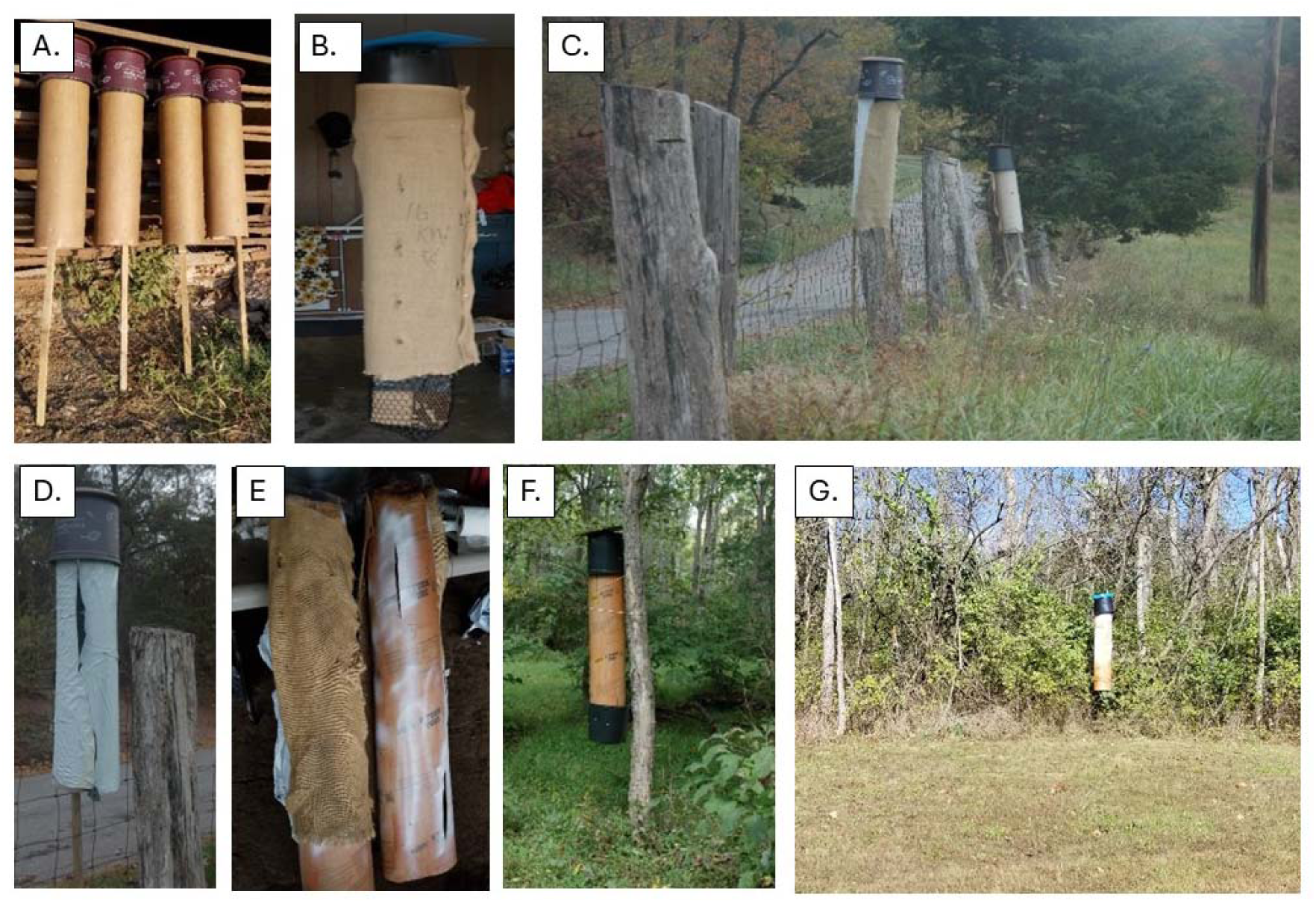
Trap #3 is presented as the “fake” or “faux” tree to liken it to dead and dying trees commonly used by overwintering BMSB in the fall and winter. The cardboard 20 cm (8”) tudes are attached to a tomato stake or similar and then covered with plastic materials like “weed beaters”. It only uses 6” or so patterns of colors of white, black and or burlap material or painted as such. These and others make the BMSB see the traps as “dead and dying” trees. A. shows the original cardboard tubes. B. shows the burlap cloth used on it. C. shows how they were placed in the field. D. shows the color with a piece of white. E. have the tubes with burlap and white plastic as well as just the tube with paint around the holes. F. shows a trap hanging outside and G. shows one of the traps that caught ∼500 BMSB along a tree line behind a large house.

### Trap 4

“Weed beater” plastic materials in black or brown were modified to vertical positions into multiple sides of trees, etc., (3-4 or so sides) as follows to provide field/forest places to hang traps numbers 1 and 2 (Fig. 13). Such built structures or enclosures could also be termed “haven”, spatial components, crags, crevasses, ledges, palisades, refuges, sanctuaries, wash outs in higher areas of height, etc. Two types of weed barriers were used - Amazon-sold “Mutual WF 200, polyethylene woven geotextile fabric in 91.4 m × 196 cm (6’5’ × 300’). The other one “Tuffin weed barrier 91.4 m × 196 cm (6’5’ × 300’) garden landscape fabric”. This was named the “Haven” methodology for using the plastic material that isn’t that expensive and its success depends on how it is setup to attract and fool the overwintering BMSB. Success largely depends on how it is organized and most importantly where. The “faux tree” traps can also be placed relative to the habitat factors motivating the movement of the BMSB trying to find overwintering places. They don’t like open spaces before the en masse flight unless a building or such is present. Both traps, e.g. #1 (Pot) and #2 (coroplast) can be used on these 2 materials.

**Fig. 13:**
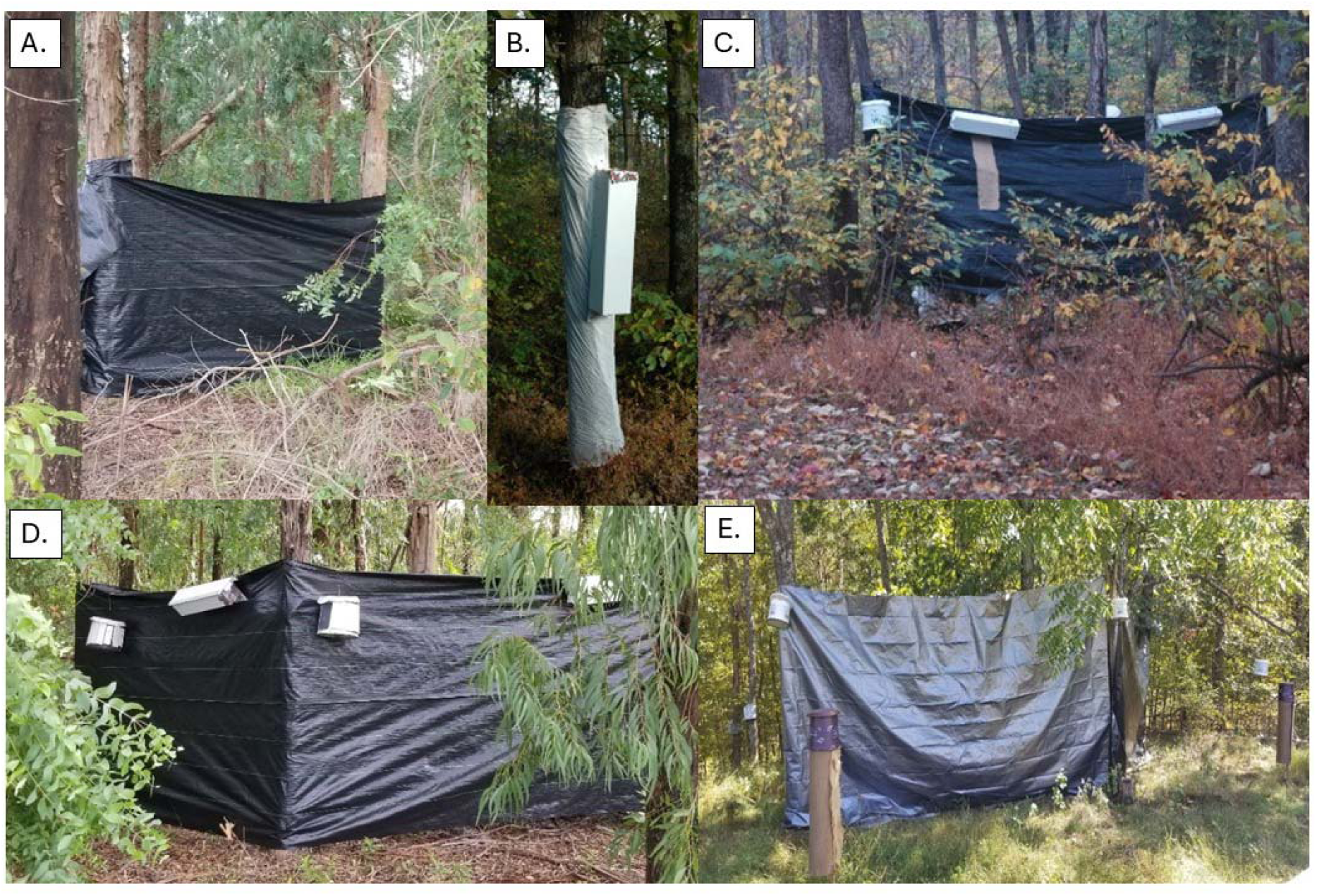
Trap #4 is developed of multiple pieces with the major portion of the plastic materials called “weed beaters” and the other 3 traps placed on or around it. These materials (“havens” for a name example) together produce and enable several behaviors of BMSB before they commit the en masse movements. There are many factors that go along and allow understanding of this invented material. A. shows the side of plastic placed around the trees to provide a high part of a Haven. B. shows a Trap #2 placed along with a tree and C. shows a trap with traps added onto the weed beater. D. and E. show how the materials might be set up in the areas when BMSB have been seen and/or attracted.

### Trap 5

Roofing materials that are placed under shingles of housing, etc. (Fig. 14) were tested and evaluated as a potentially cheaper (∼$20.00/roll) “Faux tree”. Also, cardboard boxes from Uline, Inc. can be set up with burlap and white materials to function similarly to the tubes made into “dead and dying” looking trees. With all of these “Faux trees” you build them on tomato stakes, drill entry holes (usually at the top) into them to enable enhancement of BMSB interest in added hexcel, place a plastic lid or coroplast top over them to stop water entry, stick them in the ground outdoors. Then the only use is for a few weeks before they are counted and used or moved forward for something else. This cheaper trap does work, but I prefer trap number 3.

**Fig. 14:**
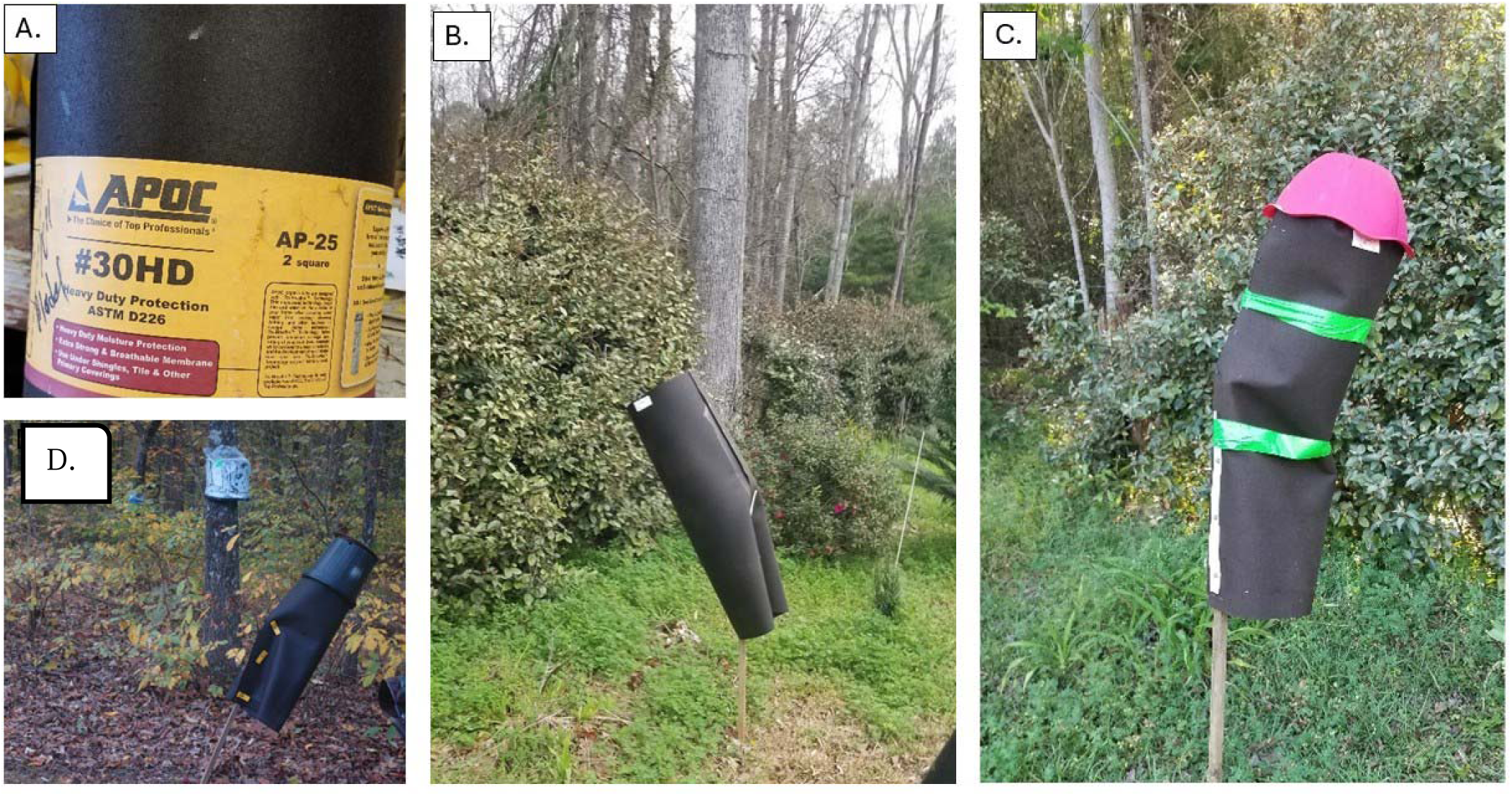
Trap #5 is made of roofing materials to present a “dead and dying” tree as with Trap #3, only it’s cheaper in costs. The trap uses the hexcel inside, has a hat on top and will work but tends to bend over. However, that can be reduced by adding some #9 wire inside in circles to hold it up. A. is the roofing material, B. shows the trap in the woods, without a hat, while C. and D. show how the finished trap can be used.

### Colors of traps and backgrounds

Insects are well known to prefer individual colors over others depending upon their behavior and needs. However, the BMSB during the overwintering time between the fall 12.7 hrs. and the spring 13.5 hrs. photoperiod initiation does not appear to be concerned about more than finding a dry, cool/cold, and safe hiding place until the spring - summer returns (Nielsen et al. 2016, 2017). This idea was tested and a good example would be to find places used in buildings, houses, barns, dead or dying trees, etc.

### Testing and evaluation of trap materials and components as developed

As the above traps show, they became the best functional and final tools, so other questions of the response of BMSB overwintering were made and tested. These included colors of the traps or things around the traps, heights off the ground and on buildings, cardinal directions of traps on buildings and other potential characteristics that might better enhance the traps. Please note that previous research on such behavior has been done by others on BMSBs when they were in the usual stages of nymphs and adults (Cambridge et al., 2017, Egrl et al., 2023) and by overwintering BMSB (Lee et al. 2014). Also, BMSB can get into something slit of 3 mm to 7mm. BMSB were tested for behavioral response to dead BMSB of both old and new and did not respond at all to dead BMSB (Mizell, observations tested).

### 2014

Fifteen m (50’) and/or 12 m (40’) high white silos 3.7 m wide (12’) (Fig. 15) were used to hang large pot like traps along the outside areas. The traps were colored white, black, yellow and silver and placed around the outside of the silo in sections with each of the 4 traps 30 cm apart. Please note that BMSB do commonly respond to silos (especially during the en masse flights), but there are no places for them to hide and overwinter on only the silo exterior. Here, five sections of plastic pots were provided and placed in the north, east, west, southeast and southwest areas on the silos. The sections were chosen due to the difference in heat that develops in certain azimuths which such “named” sections on a silo easily affect it. The results were the capture of a total of 2107 BMSB. The outcomes provided – 4 colors and 5 sections – were not significantly different by either factor. By the colors, white had 567, black 719, yellow 353 and silver 508. The sections had non-significant statistics and very variable with north 293, east 401, west 268, southeast 169 and southwest 989. The result with the highest number at the southwest side (∼210-240 degrees toward the sun), is likely due to the behavior of flying BMSB on the en masse arrival to the silos, which usually arrive in high numbers in the middle of the afternoon. The traps used in this early experiment were effective and indicated that a Pot trap could be worth preceding with testing and development.

**Fig. 15:**
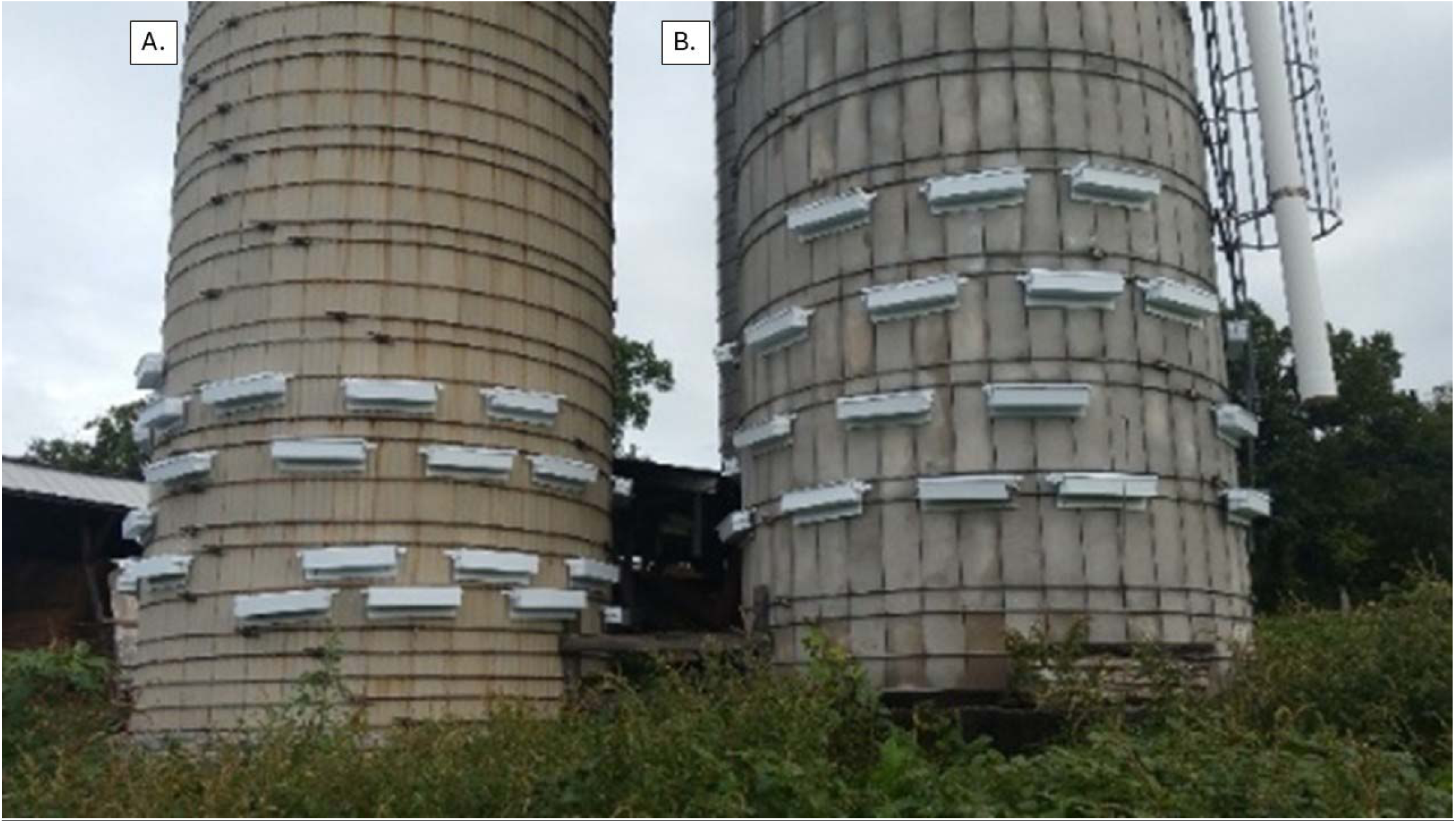
Use of traps to compare colors and heights using large silos. White, black, yellow and silver were set in the silos at 30cm apart. The traps were placed to provide comparisons of the response of BMSB to heat found at different azimuths. The results were the same, not statistically different either way for BMSB. By color, the white had 567, black 719, yellow 353 and silver 508. The response to azimuth was also variable but not significant by statistics. North had 293 while southwest had 989. A. and B. are two separate silos of 12-15 m (40-50 ft.) high used in the work with Trap #2.

### 2016 – 2017

Pot traps in 4 different colors of black, white, silver and UV mulch, and 30 cm (12”) apart were held on each corner of the cabin and placed (see pictures) so that each of one of the 4 traps and colors were mounted similarly (F-value 3.49, Pr >F = 0.098). Heights had no differences. Numbers collected were 116, 117, 95 and 72. Cardinal directions – east = 115, north = 165, south = 55 and west = 64. The least square means for cardinal directions indicated east (4.5) and west (6.4) were significantly higher than the south and west at 2.2 equal. Also, north > south (0.027) and > west (0.045).

### 2017: Evaluating the color and background colors in an open hay barn

In this experiment the color of the traps and the color of the backgrounds were tested as BMSBs moved away from the silos and other farm equipment into hay barns. White, black and silver colors were used, and the results were relatively high numbers by the traps but not statistically different. With a background color of white 621 were caught, black caught less at 491 and silver was lowest at 324. The trap colors of black caught 683, greater than silver at 460 and white at 229.

#### Use of a cabin with some open areas as well as forest trees

Sunlight, heights and spacing above ground: These factors were manipulated to try to have them used by the BMSB with the hope that they might enhance the importance and usefulness of the traps. Silos, barns, cabins, haylofts differed on some early traps and colored materials were used. Light coming from the four directions such as a wooden cabin or a brick-built home can be affected by the azimuth direction of the most sunlight at a specific time of day, the openness and placement of trees and other factors reducing surrounding light.

### Year 2017

For this experiment, the final Pot trap in 4 different colors – white, black, silver and UV mulch were used. The traps were hung on hooks over the 4 sides of the cabin at 30 cm (1’) apart (Fig. 16). Please note that UV mulch has an amazing effect on ambrosia beetles which I have discovered, part of which has been published by Conner et al. (2026) and the finished article will be available. The traps were removed to the next corner every two days along with switching the height positions of the individual traps together up and down. Thus, each trap was exposed to each of the spaces, heights, and colors for the equal amount of “convergence” time. However, the traps were cleaned and counted on each of the change days, so numbers of each trap type were gathered by direction and position. The total number of BMSB captured via this setup was 2424. The overall number of white traps was 575, black 279, silver 545 and UV mulch 628. Again, the numbers captured were high but not statistically different. The amounts of BMSB in cardinal directions were north = 1101, south = 277, east = 524 and west = 352 shown out of the 2424. The amounts out of trap positions were highest trap = 618, number 2 = 641, number 3 from the top = 567 and just above ground = 458. As indicated, the area around the cabin used was modestly covered with trees that were variable in numbers where they occurred.

**Fig. 16:**
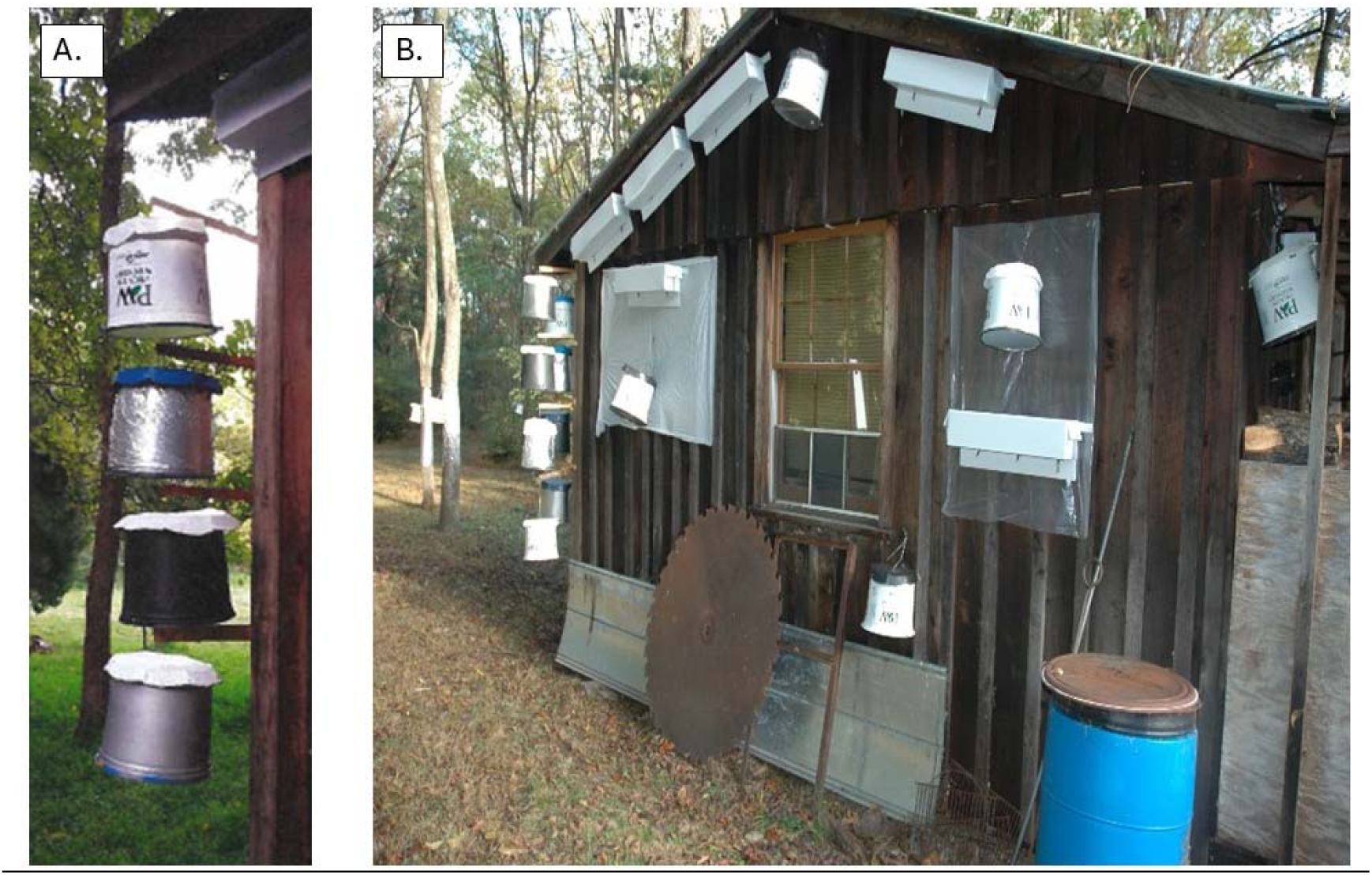
This picture shows the traps that were tested on a wooden cabin for behaviors of BMSB in response to the available traps on the outside. Factors were colors, heights and cardinal directions with traps. The information is provided in the publication. A. is the 4 different colors of traps tested. B. are the 4 traps on the edges and other traps can also be seen mounted.

### Height around a cabin area using Pot #1 and the rectangular coroplast trap #2 in a comparison of collection and height of BMSB in 2018

Cardinal direction and height around a cabin were again used to compare the 2 different traps for their cardinal direction and heights for any type of BMSB variation. On all four sides as described above, the 2 types of traps were hung side by side (Fig 17) on ladders at 0.9 m (3’), 1.8 m (6’), 2.7 m (9’). 3.6 m (3.6’). and 4.6 m (15’). The Pot trap (#1) was used normally and the coroplast trap #2 was used vertically with a lid and holes at the top. BMSB in the traps were counted after 3 weeks. Results for direction comparison show that the 187 BMSB collected in the east were greater than the 96 on the south which was greater than north 63 and 18 from the west. For height the comparison of the two types of traps indicated that Pot #1 caught more than #2, the coroplast trap. The collection was arranged by Pot 2.7 m (9’) > 3.6 m (12’) > 4.6 m (15’) > 1.8 m (6’0) > 0.9 m (3’). The actual numbers are Pot #1 at different heights 119, 90, 63, 54, and 38. The coroplast trap results were 3.6 m (12’) = 2.7 m (9’) = 4.6 m (15’) > 1.8 m (6’) > 0.9 m (3’). The actual numbers were 72, 68, 64, 48, 28 for #2. There were several similar experiments that were tested to help develop important and effective traps that could be used to work against BMSB. However, the overwintering BMSB overall showed very little interest in anything other than to find a dry place that could be used as a hiding place for the winter out of the sunlight, without water, with more than 1 BMSB per hiding spot and some addition perhaps of other unknown factors.

**Fig. 17:**
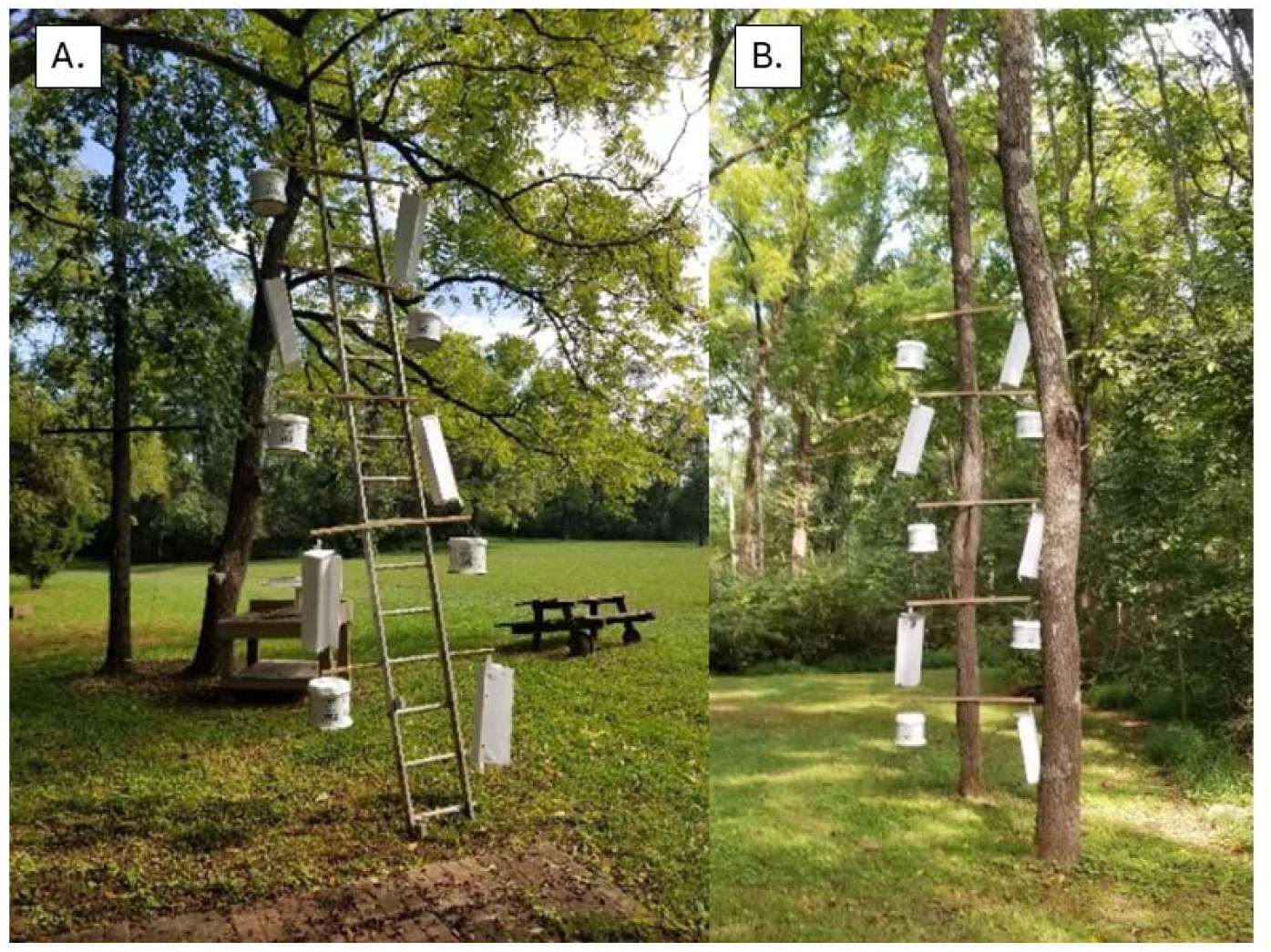
This picture demonstrates the comparison of Trap #1 and a vertically hung Trap #2 for the use of cardinal directions and heights. A. and B. both show the ways the ladders and trees were used. The results indicated a small difference in the cardinal directions use and that Trap #1 caught a few more BMSB than Trap #2 vertically.

## Discussion

Completing this paper targeting BMSB overwintering behavior and invented traps occurred in early 2026 and by now there are a high number of publications from all over the world since the BMSB spread to most places where agriculture and related areas occur. Lee et al. (2013) and their references reviewed the management of BMSD in Korea and reported many of the same factors as in the U.S. and Europe. Wallner et al. (2014) evaluated the use of 48 landscape/land use variables looking at urban, wetland, forest, agriculture and railroads in New Jersey. Leskey and Nielson (2018) summarized the information of subject matter found from North America and Europe including impact and management. Bakken et al. (2015) evaluated the occurrence of BMSB along forest edges as well as suburban settings in North Carolina and Virginia. They also investigated soybeans over 3-4 years. No managed plants or the soybeans in the coastal plain areas were low, but much higher in the piedmont and mountain regions. A number of plants were recognized to be preferred hosts: tree of haven, catalpa, yellowwood, paulownia cherry, walnut, grapes and redbud. This information was helpful in predicting attractive areas for BMSB population increases. Lee and Leskey (2015) also found that after emergence and overwintering sites in spring, numbers reached the highest in June. However, in the field, flight direction was opposite to the sunlight positions, flight direction was opposite to sun position in early PM.

Morrison et al. (2017) found that BMSB responded to pheromone aided traps after critical 13.5 h photoperiod in the spring. White traps were found in 8-20% where locations found it was not effective for overwintering management indoors. Hancock et al. (2019) used volunteer gathers for data collection. They looked at home exteriors during the peak dispersal as well as home locations with color and structural materials. Adults were counted more on the white and white-tan homes and grays. More adults were counted on the north and east walls and on homes with wood, cement and stone. There were also more adults in rural landscape homes and more on darker ones of natural materials. These types of rural homes were prone to large nuisance infestations of overwintering BMSB. Cianci et al. (2021) investigated the outside overwintering BMSB in Ontario, Canada. All died but ones in shelters or other similar things lived. Such BMSB are chill susceptible. BMSB are dependent in Ontario on built structures and other microhabitat’s available to overwintering BMSB.

The main observations were that the BMSB are a nuisance as overwintering adults. BMSB, unfortunately, also is a polyphagous species with a strong dispersal capacity, has a high production rate and can spread into most regions often as/after the “overwintering” finishes in spring. Furthermore, the occurrence of overwintering BMSB and the remaining potential to become important pests stands still as potential large losses across many food production areas due to the lack of acceptable management tools.

Everything that I observed and worked with BMSB on the farm occurred during September - October relative to setting out and then taking down and counting the BMSB. As part of the years of work, I invented quite a few traps to test but am only presenting 4 traps also with a “methodology” that goes with them to present an attractive “Haven” (a place of safety or refuge), but also may be a cave, enclosure, cavity, harbor, or sanctuary, etc., that attract large numbers of BMSB and on which the traps can be placed in the landscape to capture the insects as they pursue overwintering site(s).

Here remember that one of the important pieces with any trap is the material inside, the hexcel is used inside the traps, and is the key material used by overwintering adult BMSB! The 4 traps include #1 that is a plastic pot used for nursery production but works by hanging it on buildings and elsewhere. The Pot trap can be placed around buildings, etc., including the Haven or something similar, where BMSB are moving and looking for acceptable areas. I have caught anywhere from 0-100’s to ∼4,500 BMSB in these individual traps.

The second trap made of rectangular coroplast that is pressed into shape 0.6 m (∼2’) long by 13 cm by 15-20 cm (6”) rectangular square that you can place on the walls of buildings and elsewhere. BMSB go in the traps and stay until spring unless there is some unusually warm weather in which case they come out and mill around then usually return. The traps placed on the walls (and Haven) are very efficient if placed 46 × 61 cm (18-24”) apart.

The third trap, a tube shape fake or “faux” tree mounted on a tomato stake, is placed around buildings, etc. The “faux tree” success also depends on where it’s placed all relative to the habitat factors motivating the movement of the BMSB trying to find overwintering sites The tudes were also tested using ∼white plastic materials, burlap and burlap paint pieces 13 cm (5”) as smaller additions to the faux trees to improve the added BMSB attraction of the tube traps as dead or dying trees.

The fourth trap is a grouping of 1, 2 or all of the #1, #2, and #3 traps mentioned and added along with the black or brown “weed beater” polyethylene materials used normally on the ground for control of weeds. The use of the trap basically enables making unusual “buildings” that can be placed most anywhere for evaluations of attractions. The “Haven methodology” isn’t that expensive and its success largely depends on how easily it is set up off the ground 1.8 m (6’6”) to form 3-4 sides stapled to trees or poles, etc., to attract moving BMSB. Traps #1 and #2 can also be placed on it to attract the BMSB and #3 can be set up in the areas close by the #4 to capture and or try to find where overwintering BMSB are present or moving about. Moving and wandering of BMSB is quite common and a term for what is looked for can be called, e.g., “convergence”. That is the place where all kinds of factors are found most anywhere and made from many things!

As other researchers have found in a final list of activity evaluation, overwintering BMSB do not like open spaces, high heat and direct light, nor wet areas, but do like buildings and the like (trees of various type depending upon location), locations of old garbage metal, cars, trucks, farm related stuff such as silos and the like of most any kind that meets the last sentence list and both before and after the en masse flight occurs (see below).

### Overwintering behavior, observations at the farm

BMSB that are alive in August when the day length reaches around 12.7 hours/day the insects go into dormancy and begin to move through the landscape (Nielson et al. 2016, Morrison et al. 2017). On the farm in early September, it appears that males and smaller BMSB are the ones observed moving about first. This occurred weeks to a month or so after the daylength change where I was working and there is a lot of flying by these insects in the woods and other vegetation habitats. BMSB will fly around buildings and trees, etc., (my observation) while looking for dead or dying hardwood trees which they go into but at low numbers/tree (Acebes-Doira et al. 2015). Again, they don’t like to fly a lot in the open unless around buildings and such, they do not like plain open areas and prefer something around 50% vegetation in the flight areas. This is how the BMSB work (my observation on site) and it is how the “weed beater” was setup to use. The actual types liked by overwintering BMSB are very similar to the normal nymph and adults moving around in the 12.7 P1 occurrence that will be the 13.5h population in the following spring (Cira et al. 2016).

When the weather gets cooler (this is similar in BMSB as it is in *Harmonia axyridis* Pallas (HA) but not 100%) (Mizell, personnel information), HA requires more and colder weather to switch, and about temperatures below 55-60 (the lower the better), they start landing on houses and other “havens” that fit their visual, etc., requirements (Mizell, personal information). Note: as stated earlier, the usual testing of trap colors, trap heights, etc., of BMSB was done but didn’t really find anything very important. After about 3-5 days or more (if it warms up) of the cooler weather, BMSB will start flying in an en masse cloud often in the open and at higher numbers from an azimuth angle of ∼210-240 degrees (the southwest at ∼2:00 pm) or so and go to large building and other things in the skyway such as silos, houses, barns, etc., especially ones facing south – southwest. Once landing they do not stay on the silo because there is no place for them to hide on a concrete silo. But again, if you placed #1 or #2 type traps on the silos, the BMSB will use them readily. They do go into the surrounding farm buildings, houses, silage augers, etc., anything dry, hidden and close, that has places to hide in. Also, as indicated, hexcel is inside the traps and highly attractive and effective. It is also easy to use for counting and/or maintaining live BMSB from the traps with overwintering BMSB for your science projects. They will wind up in all the buildings surrounding the silo for at least half a mile or more if the land is both “open” and “closed” (corridor, barrier, matrix, and convergence, see below) that enable covered areas and can be seen from the higher silo. As do HA, overwintering BMSB enter the houses if they can, go through its openings and come out later in the inside of the buildings.

We know that BMSB, as do most stink bug species, have a strong “edge effect” in their movements through habitats and corridors during the summer (Rice et al. 2017). In the summer, this could/would be the landscape ecology level behaviors known as corridors, barriers, and matrix. This type of information was involved in understanding fragmented landscapes in the southeast like a few other places some time ago aimed at improving the numbers of habitats and their insects and other organisms of interest (Forman 1995, Tewksbury et al. 2002, Haddad et al. 2003, Fried et al. 2005). Matrix here means the vegetation parts that are graded by density and probability of usage; more often not used, too thick in vegetations is one example why. And yes, convergence of things involved with BMSB work also are similar. Please note that the information and methods involved in the more interesting understanding of how BMSB respond to weather factors and can chose the proper times to fly in en masse flights will be provided in the “Supplemental” section. If we can get the AI approach done?

At this point we turn back to the target here, BMSB. While corridors are wanted to be used to enhance positive insects and other related animals, here we are trying to target BMSB when they are trying to protect themselves for the winter and the rest of the time used over the winter to spring. What must be understood and to look for is the corridors and barriers so you can strategically place the Havens with the traps in acceptable corridors and convergences with them. On a farm with lots of areas where there are changes in the landscape by variations of roads, fencing, forests, etc., various other things such as unused old cars and trucks, dirt roads, fence lines, tree lines, woods, etc., it is termed “convergence” (by the author) and can be used as a clear idea to work with (Fig. 18).

**Fig. 18:**
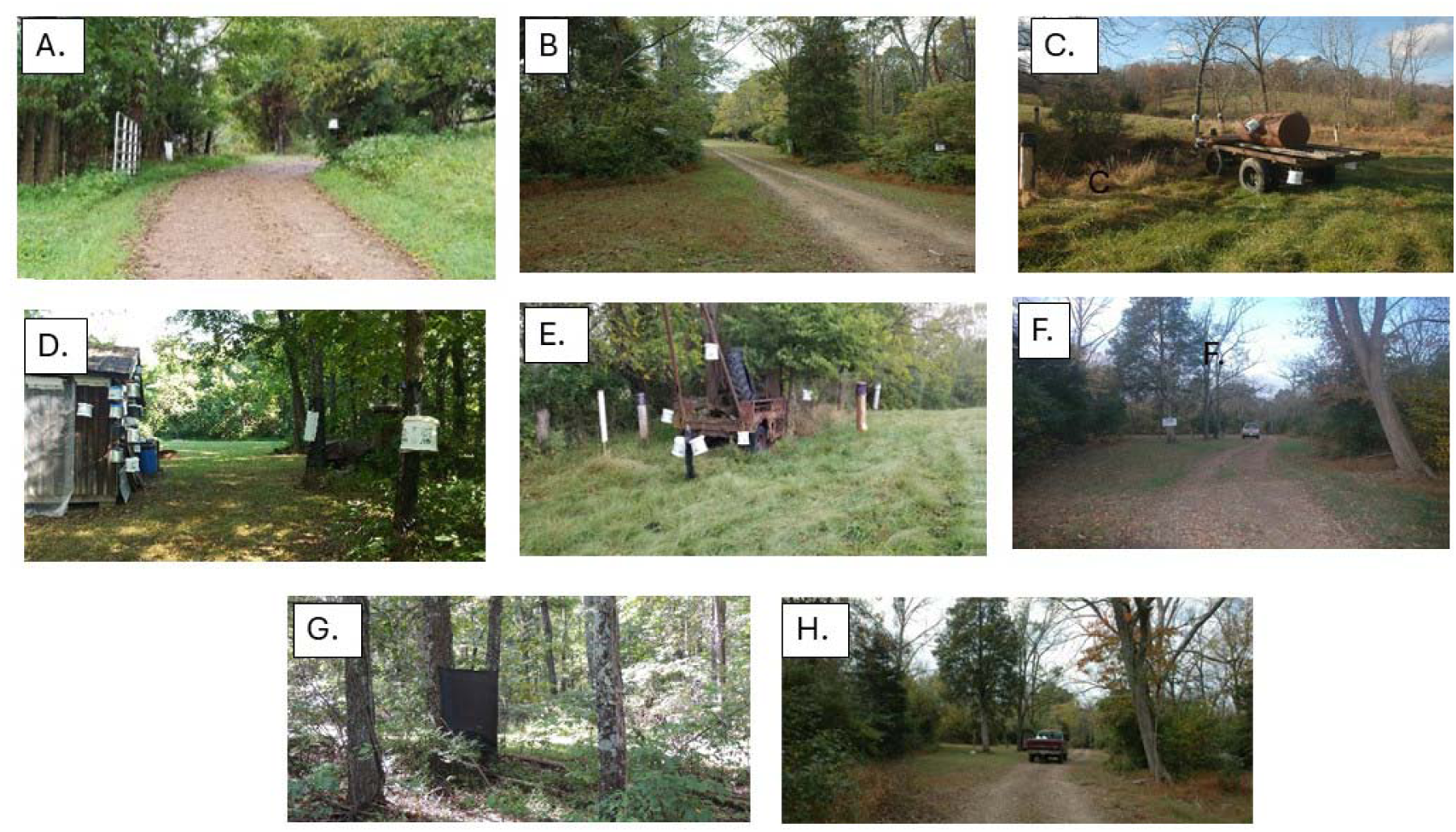
These pictures show the type of “things” that are found, made, turned into junk and the like from manmade materials. They also show the “convergence” that happens when all kinds of “things” occur such as trees, roads, mistakes, and whatever due to change over time, etc. Landscape ecology management scientists some time ago talked of coroners, barriers and matrices as ways to understand and improve places that increase insect populations and such. Here with BMSB we wish to do the opposite. For the BMSB, we have all of these factors and should try to manage better in winter. A. shows the dirt road entry point and B. shows the other side as it looks when entered that way. C. shows a pile of junk placed in a farm field to attract BMSB which it did not. D. is an area around the south side of the cabin where BMSB do fly around. E. is a large site past A. which is an old place with a junked truck. F. is a somewhat open area further along the trail from the truck. G. is a piece of weed beaters set up close to the south side away from the cabin. H. holds a truck having passed through all of the A-G areas mentioned as the truck is moving toward the cabin from the east side. This area also shows a lot of “convergence” and BMSB were observed trapped in all the places mentioned and shown here.

As a testing example, large numbers of various materials in all sizes and shapes were placed over the farm as potential attractions along with convergences and habitats as well as wind, water, vegetation, junk, etc., to the BMSB during the overwintering behavior occurrence in 2018. This material was placed in 12 areas to find trap captures in response to placement and type. The mixtures of the 12 locations were placed in different parts of the farm within different types and sizes of vegetation, trees, open areas, etc., as well as “convergences”, corridors, barriers and matrix (Tweksbury et al. 2002). Given the invention of the 3 new traps, the “weed beater” (Haven), the roofing material and the behavior observed and somewhat described above, an idea was raised of trying to develop “Havens” that could be placed out in the habitat with the traps on them as seen fit relative to the population of insects, the habitat around them, and most importantly its structure (meaning forest, usual vegetation types, etc.). “Convergence” areas occur over time and are generally made by available stuff real, broken or stashed, and their use can help attract BMSB during overwintering and the addition of the traps hung in the areas are very useful. As indicated above these were included on the farm, are old wagons, tractors, other types of equipment, trees and poles along with addition of the invented traps and Havens. Cardinal directions around materials make up some importance to consider as light and health are mainly on the sides of the materials that receive more sunlight and heat not usually found in the north and east directions.

What does this mean after the availability of the mixtures? What was learned and tested from this effort deserves about a 80-90% rating. I know what is needed but it can be hard to come up with key areas that the BMSB will be attracted to and moving through to choose the most effective place to set up the Haven(s) with traps. Such things as convergences do occur and can be figured out for use in known areas and types. For example, foresters evaluate sites according to the available forest production needs at present or later (Elledge and Barlow 2018). As a side comment, I’ve developed trap cropping plots very effectively for *Euschistus* spp. along with another trap (yellow pyramid) that I invented (patent was granted in late 1990’s) and these Haven ideas are very similar in analytics, evaluation and development (Grabarczyk et al. 2022).

As a result, my statements so far need an estimated percent of how sure that I am of what I have made and written. The traps #1 and #2 get 100% rating, if the BMSB are near them, they will enter them and stay at some relatively high numbers. The #3 faux trees get a 80-100% support that is dependent on where they are placed, the population of the BMSB, and characteristics of the habitat. However, similar activity can be found if pathways or “corridors” occur along areas with heavier vegetation.

### The outcome is this

With all the items I have developed and discussed here, I believe that they can be put together and used to manage a high population of overwintering BMSB as follows to complete one or both of two management strategies starting after 12.7 hours early in the year of P1. The invented traps #1, #2, #3 and #4 (“weed beater” and traps) are ready to use properly. It’s the “convergence” materials, and as indicated corridors, barriers and matrix that must be managed correctly and effectively. There is no “maybe” here because of corridors, barriers and matrix, its monitoring, handling and using to win.

Let’s say we have an organic vegetable grower with a few acres that each spring and early summer (after 12.7) are bombarded by BMSB from overwintering buildings, surrounding habitats, etc. By using the “Weed Beater” = Haven traps, there is now available these new, special tools that can be developed and used to ask and answer a number of key questions about the spatial-temporal movements of overwintering BMSB that will allow a much better understanding of the details of their behavior on “when”, “where” and “how” the BMSB set up for the winter and the beginning of the P1 group (13.5 h) for spring/summer just beginning. Any person in charge of the following suggestions must first understand the phenology and physiological characteristics of their local areas throughout the year. The populations of BMSB that occur at the end of the season and then become the overwintering populations should be developed and understood per year. As I have suggested adding in the combinations gained by deploying the “Havens” (use what works for you) in key places, the farmer should be able to: **1.** Use them without losing a lot of money to set up the traps and harvest them relative to the landscape/farmscapes areas. **2.** Using “the” or your “Haven”, find out whether overwintering BMSB do or don’t move around in the fall in relation to the suitable places they might use for movement and useful “convergences” present man-made or else build up some new tools if you don’t have them. **3.** Delay the damage of the new BMSB pest to be 12.7 h and increasing into the summer to vegetables, etc., by lowering the overwintering numbers using the traps in the fall and winter with the best time and places. **4.** If necessary, add the use of a minimum amount of insecticide that is still needed. **5.** Evaluate, analyze and change as needed, then repeat until a good understanding of BMSB management is accomplished within the areas working in, etc. That would include “convergence” materials available (natural or built, e.g., “weed beater”), any of the factors on the lists here and elsewhere of such related to management of BMSB summer and winter. Use the tools properly as suggested and remember the background materials provided above when needed. Depending upon where one lives, the time from fall to spring available to control the pests in most places is available for months.

### Other things observed and evaluated

Tests were conducted with traps to develop what type of available additions would work to increase the number of BMSB within various areas as found and manipulated over the Mizell farm. After over 75 years of family farming there were plenty of factors that could be used such as the various types of farmland and forest types (e.g., open in grass to areas with 25, 50, 75% of different types of forests), and “convergences” with lots of old trees and stumps, lots of broken machinery, old and new and moveable. Also, there were 3 houses and a cabin with grass around and lots of other smaller buildings around them to test and manipulate with the 4 types of traps. The available and buildable haven areas could be placed in numerous different types of closed, open and convergent areas with different vegetations (amount, types, sizes, thicknesses, etc.) for comparison, etc. Such things are available, easily makable and can be used.

### Auto-diffuser

**Novel use** of collected diapausing BMSB as an auto-diffuser of pathogens and/or growth regulators to enhance the spread and efficacy on overwintering BMSB for a releasing epidemic. Find out where the overwintering BMSB are and treat them early, late or” best” if you have it in the spring across the areas that are the pest problems occurring. Or collect them in spring and treat them with such companies in the industry with bacteria or fungi that can spread among the BMSB populations across the crops, etc.

## Conclusions

### Brown Marmorated Stink Bug Overwintering Behavior, Trapping and Monitoring

At this point I conclude that an overwintering trap can be used to:

1. Quantify and determine the annual end of season population levels on an areawide basis,
2. Estimate the overwintering beginning populations locally and areawide,
3. Collect data on the landscape factors you see and how BMSB use them when and when not. Do not forget to take good information on the developed and often natural “convergence” positions of the BMSB over time in your area(s),
4. Indicate the annual BMSB phenology (and to help predict it – see #4) in relation to overwintering en masse flights and emergence,
5. Define, evaluate and determine amount of BMSB and where they are prior to en masse movement and afterwards,
6. Be used to develop temperature driven predictive models to estimate various population dynamic parameters (maybe combine it) with data from *Harmonia axyridis* sampling),
7. Use traps, etc., for several other experiments (or facilitate such) to further understand BMSB behavior, ecology, etc., toward the implementation of improved IPM strategies and tactics. In other words, the public’s money gets well spent!
8. Get data on the overwintering en masse flight factors that will be the real things needed to do better control of the overwintering BMSB.
9. What is the best way to stop the BMSB from effectively moving and landing into new summer conditions and functioning as they normally do?
10. Are there any other insect species that might behave similarly enough to be trapped using similar traps and materials as used for BMSB?
11. University of Florida “AI” results are coming and perhaps show how that BMSB and HA decide on when they fly en masse. If so the results could be very useful for IPM needs.

## Competing interests

None

## Acknowledgments

This research was partially supported by the USDA, National Institute of Food and Agriculture, Specialty Crop Research Initiative under award number 2016-51181-25409. I thank Dr. Anne Nielson from Rutgers University for leading and supporting the research. I recognize and thank my wife, my brother, my two sisters and their husbands for helping at the farm during the work on this project.

I thank the following for reviewing an early issue of the manuscript:

## Supplemental Information

**Fig. 1:**
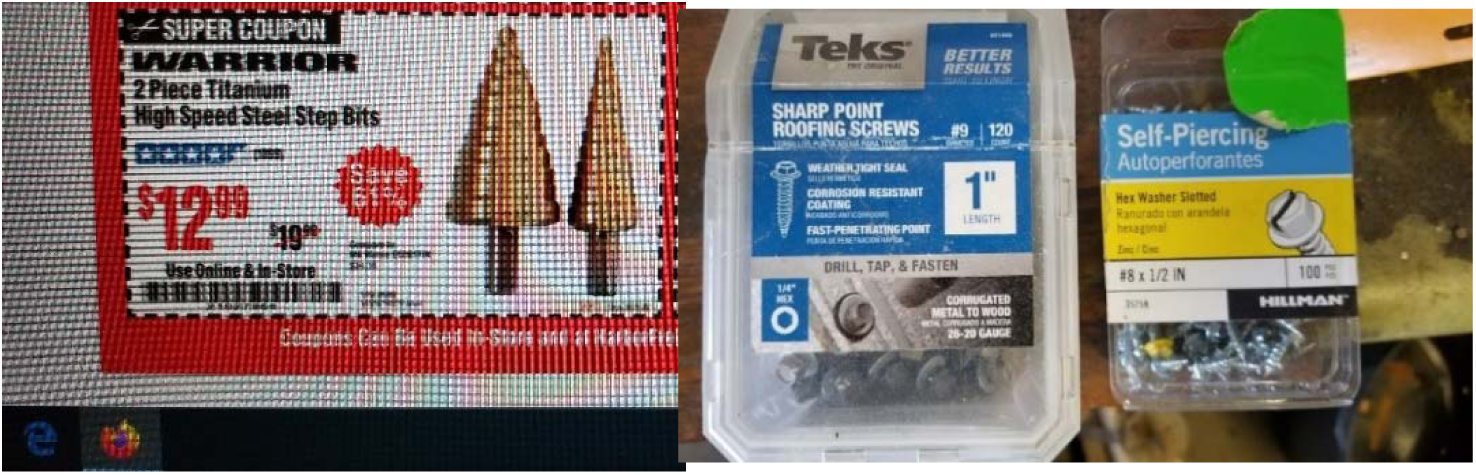
These are tools used to build the traps easily, use them, then to take apart and put away until next year.

### First BMSB found in Florida

The first BMSB reported in Florida was by Penca et al. (2018) in Lake County, FL in peach orchards in 2016. They were found in 2017 in Poke County, FL and the eggs and nymphs were found in Lake County by 2018. During these years it was well known that BMSB were “hitch hiking” on a number of vehicles that wound up in Florida or other southern states from other northern invasive states. Here is some unusual data gathered earlier in north Florida around the times when BMSB showed up in U.S.

### Movement of insects in recreational vehicles (RV): unknown spreads of pests too late!

In 1995 I held the first position 50% of my time of being the first head IPM agent at the University of Florida-IFAS. I lived in north Florida as a Professor of Entomology working with fruits, nuts, forests and nursery pests starting in 1982. The north Florida agricultural science area was ∼25 miles from Tallahassee as the Monticello-AREC. That area is almost 2 hours from the University of Florida in Gainesville and now serves as the county Extension office. There are 20+ locations with agricultural scientists that serve similar jobs helping the public reduce the effect of pests of all kinds as well as many other related problems related to agriculture. My job was to work with other scientists to find support funds, talk to the growers and address ideas such as reducing insecticide use, more natural enemy use, and the like. Most of the agricultural science places were and are still in central and south Florida which caused me to drive long distances across the state to serve my job.

There are three major dual highways, “I” is for interstate, roads that occur in north Florida. Beginning from the east and the Atlantic Ocean, the I-95 is in the east and runs from the far north (example Washington D.C.) to south Florida. The I-75 is west of I-95 about 150 miles also running parallel in the same directions. The third road is I-10 which runs from far west Texas through the rest of the east to Atlantic Ocean at Jacksonville. In north Florida are the towns of Live Oak, Monticello (where I live), Tallahassee, Panama City, and Pensacola, FL. Great roads, but long and tiresome rides to get there, which brought me to the current insect related story and to support (Fig. 2).

As every traveler knows, driving is tiresome and repeated driving often is worse. To avoid the “tiresome” problem, I came up with an insect-related idea: ask some questions and see what the “road” might provide of interest and at least keep one awake. On I-75 there is ∼45 min. between leaving the I-10 for the I-75 and driving to Gainesville, FL and the University of Florida. So, I decided to use that 45 min. to do, an experiment to examine RVs (recreational vehicles) that I observed often “going and coming” “arriving and leaving” along with the usual semi-tractor trailers. The RVs used were of different types but ones that were big enough and the type that would be able to be associated with insects. Back in those days RVs were very common in large numbers on highways. For you current entomologists and friends, the layout of the experiment and data gathering were as follows.

On each trip “down and back” from home, I used a location on the same places of road indicating good road, some distance away from the recreational areas, and the open clearness enough to see the RVs on the opposite side of my vehicle going the opposite direction. Ten miles were divided each time into 10 1-mile sections in which to count the number of RVs per mile. The mile distance was determined by the car’s odometer reading at a common speed for the 10 miles. During returning home, the 10 miles were counted again on the opposite side to provide the number of RVs going both directions for that date. A notebook was used to copy the numbers/date/mile/by directions. Over a period of time of about 2 years off and on the numbers were collected as indicated and did include the main use of I-75 and some with I-10. The RVs were much higher in number with I-75 than I-10. The collected data was analyzed by forming a means of each of the days with 10 miles and separated by the road used. The means are presented by the year, day and roads used as the mean number of RVs per mile.

Given the results and the movements found about the overwintering behavior of brown marmorated stink bugs, I present this figure and data as an interesting way to work on insect movement. Given the time of the data gathering in 1995 or so, and the large number of RVs present then, it is highly likely that overwintering BMSB were first brought to the south by RVs. Furthermore, similar activities have been and likely will continue moving from north to south or elsewhere using RVs and related automobiles, etc. The results in Fig. 2 are as follows:

**Fig. 2:**
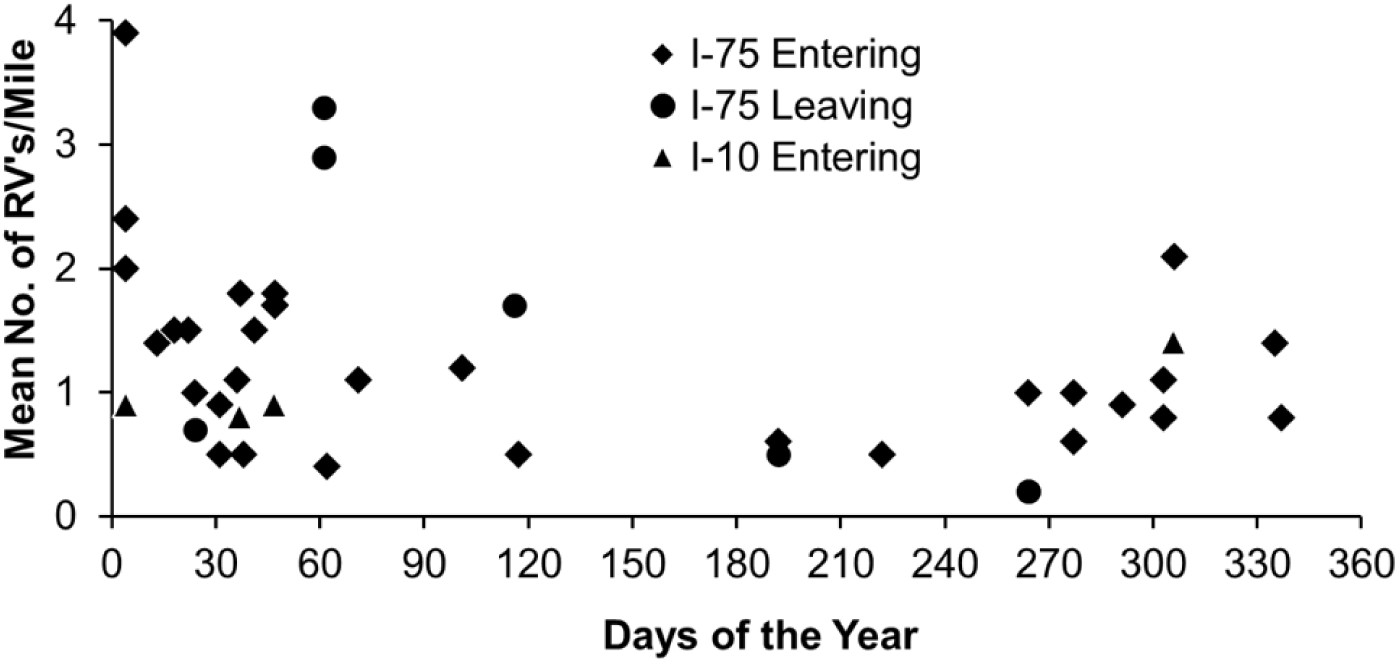
The entering and leaving times and numbers when RVs traveled on the international highways in Florida in 1995-96. The results indicate that RVs moved during the fall and winter.

---

**Supplemental Fig 1:**
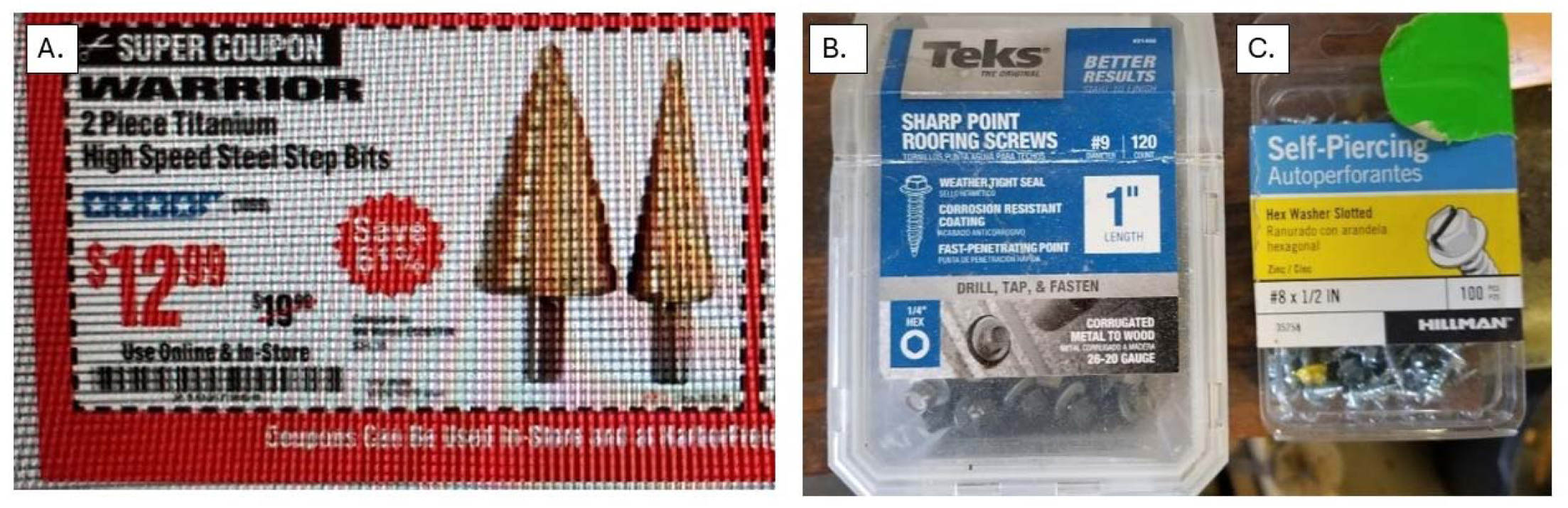
Already in the place with writeup.

## Notes

### Competing Interest Statement

The authors have declared no competing interest.

