## Supplemental for "Novel, visual-attracting traps, accompanying technology, and related behavior of overwintering brown marmorated stink bugs (*Halyomorpha halys* Stal)"

**How to Use the Mizell Traps Invented for Brown Marmorated Stink Bug (BMSB)**

**During the Occurrence of Overwintering Behavior,**

**Special Explanations**

**Dr. Russell F. Mizell, III, Courtesy Professor, University of Florida,**

**NFREC-Quincy, Quincy, Florida**

**The Pot trap (#1)**: Note: all of these traps and their pieces are important from an engineering perspective and have been tested in several ways, including sizes, colors, heights, etc. I did not discover or observe results by BMSB of some factors like colors, etc., being that important. The important things: 1. it is the angular mount of the holes (entry holes on the chosen pot). Why, because the pot 11.35L (3.00 gal) from PW (Proven Winners LLC, Sycamore, IL, 60178, 1-877-865-5818, www.provenwinners.com) is turned upside down and the bottom is the entry at top with a handmade lid on it that doesn’t close the holes since they are angled enough to stay open and without water entry on the hexcel. You can use a small screw between the lid and the Pot to hold them together better. 2. The bottom is easy to put together or take apart with 2 special nails (powder drive pins) and there are entry holes there also. 3. The hexcel is the “magic” material for all traps because the BMSB, once they enter (based on my observations), never appeared to leave even if the temperature warmed up enough for them to crawl outside the traps.

The “r**ectangular coroplast” trap (#2)** go on the eaves (flat orientation) of houses, etc. In the first test on silos, it was very effective during the en-masse flight and picked up a high percentage of the BMSB when they landed. Place them at 46 – 61 cm (18-24”) apart so that the traps are not close and do not overlap the attraction of the BMSB, and you’ll have more area available to cover with fewer traps. You can also turn the coroplast trap into a vertical orientation (see colored pictures) by adding a piece of coroplast over the top as a lid and down the side, too, to keep the rain out. Change the entry holes at the top under the “lid”. Changing the inside hexcel is not necessary. One important thing to remember is that the inside of the coroplast trap needs to have some small pieces (spacer(s) under the open side so that the BMSB have access and can enter the holes that might otherwise close when the larger pieces (4 x 4 x 24”) of hexcel touch the wall without a spacer. Also, you will want to slip a piece or two of a small twig in between the flap at the bottom (saves the use of holes that are not needed) and where it meets the box bottom. That will open an entry hole for the BMSB entering by the flap and then removing it when they are gathered and the “twigs” removed from the coroplast. The coroplast rectangle trap can be used anywhere, tree trunks, buildings, fake buildings, you name it. It is just a little harder to use than the Pot trap but just as good. Both are easy to count when you set up simple ways of shaking the BMSB out of the traps on to a plastic container with a cup and a collector below it. Put plastic peanuts inside so the BMSB can fall and stay alive into the collector.

The “**Faux tree”** **(#3)** trap is self-explanatory once you know the above, but with burlap or similar colored paint on it. It can also be partially coated with other plastic, clear, white or black.  It can be hung in a tree or on a tomato stake, next to buildings along corridors, fences, etc.  Brush them when new with Thompson water seal or Flex seal (or others) and use a lid out of a plastic food plate or similar and hang hexcel inside. You can add holes around the top under the lid and elsewhere. These tubes are used to make concrete footers and can be expensive. Square boxes of similar sizes will also work (available from Uline, Inc.). Twenty cm (8”) wide is big enough. You can also use the black roofing material **(Trap #5)** to make the tube (same as Trap #3) but need to have some wire bracing inside or they will go flat. Also, they are harder to store.

Finally, **“Weed beater” (#4)** the landscape fabric “material”, is plastic and available in a number of types in rolls (details are provided in the other parts of the publication) most of which are 1.98 m (6.5’) wide used to make a “haven”, crag, building or whatever you decide the BMSB view it as, placed around 3-4 trees (3-4 placed multiple dimensionally) and then once the material is attached (use it sideways to do the wrapping as needed) at the top hang on the other traps as available.  You put these “fake buildings” or whatever you and the BMSB think they are, in the “corridors” the insects fly through or toward places to or via “convergences” to overwinter. This trap setup is useful to help better understand the behavior of BMSB in higher trapping numbers as well as better explaining why and what is responsible for the location factors that are used. Following such as ideas in trap cropping (Mizell et al. 2008) and landscape ecology, components (convergences) are the areas that will improve their interception levels. Knowing what and where the BMSB are passing through is key to picking the best areas to place the materials.

**Note:** there are a number of pictures provided in the Figures section that go with the main part of the publication and that indicate where and how the traps and their materials were developed, evaluated and used as well as some related supplemental material.
